# A genome-wide survey reveals a diverse array of enhancers coordinate the *Drosophila* innate immune response

**DOI:** 10.1101/2025.09.24.678314

**Authors:** Lianne B. Cohen, Tamara Hadzic, Caitlin Sauer, Julia R. Gibbs, Zeba Wunderlich

## Abstract

To defend against microbes, animals regulate a complex immune response. The *Drosophila* innate immune system deploys a large transcriptional induction of signaling proteins, anti-microbial effectors, and other critical immune factors. This transcriptional response is encoded in enhancers, *cis*-regulatory sequences that modulate gene expression by binding transcription factors (TFs). While enhancers and transcription factor binding sites (TFBS) have been identified for several immune responsive genes in *Drosophila*, most enhancers that regulate immune-induced genes are unknown. By identifying enhancers, we can understand how their composition controls expression and contributes to infection outcome. We employed STARR-seq (Self Transcribing Active Regulatory-Region sequencing) in a hemocyte-like cell line to identify immune-specific enhancers across the *D. melanogaster* genome and performed ATAC-seq in hemocytes extracted from adult flies to assess the chromatin state of these enhancers before and after immune stimulus. We identified thousands of enhancers responsive to IMD stimulation, one of the two primary immune signaling pathways in *Drosophila*. As expected, immune enhancers are enriched for motifs of Relish, an NF-κB factor, and Kay/Jra, a bZip heterodimer pair, involved in the Imd and JNK pathways respectively, compared to enhancers active in unstimulated cells. However, when grouping enhancers by their target gene’s expression timing or functional role or by the enhancers’ chromatin accessibility pre- or post-stimulus, different groups of TFBS motifs are enriched, suggesting distinct regulatory logic for different parts of the immune response. Identification and characterization of the diverse array of enhancers that regulate the innate immune response expands our understanding of how animals fight infections.

## Introduction

When encountering pathogenic microbes, animal hosts must regulate an effective immune response to survive infections. The fruit fly *Drosophila melanogaster* relies on its innate immune system to defend against invading pathogens without the aid of an adaptive immune system (Lemaitre and Hoffmann 2007; Buchon et al. 2014). From the fat body, a liver-like organ, and hemocytes, *Drosophila* blood cells, flies produce a large transcriptional response, inducing over 1,000 genes upon challenge with a diverse set of bacteria and fungi (De Gregorio et al. 2001; Troha et al. 2018; Ramirez-Corona et al. 2021). The resulting proteins carry out a variety of functions, including killing pathogens, relaying signals within and between cells, redistributing metabolic resources, and balancing the costs and the benefits of a persistent immune response.

The immune transcriptional response in flies is regulated by several intertwined pathways that modulate the activity of key transcription factors (TFs). The Imd pathway responds to DAP-type peptidoglycan, predominantly from gram-negative bacteria, and activates the NF-κB TF Relish. The Toll pathway also activates an NF-κB TF, Dif, and is activated by fungi and Lys-type peptidoglycan from gram-positive bacteria (Valanne et al. 2011). Though often presented as independent pathways, the pathways interact in several ways – some genes can be induced by either pathway, heterodimers between Dif and Relish have been reported, and some immune stimuli appear to stimulate both pathways to some extent (Tanji et al. 2007, 2010; Troha et al. 2018; Valanne et al. 2010). JNK signaling can be triggered by Imd stimulation via Tak1, a TGFβ-associated kinase, activating the Kay/Jra heterodimer, also referred to as AP-1 (Tafesh-Edwards and Eleftherianos 2020). Mutation of key genes in each of these pathways results in loss of immune-responsive transcription and can lead to decreased infection survival (De Gregorio et al. 2002; Bond and Foley 2009; Brun et al. 2006). Detailed RNA-seq experiments have found these pathways can orchestrate temporal and pathogen specificity within the immune response (Troha et al. 2018; Schlamp et al. 2021).

Transcriptional programs are encoded in enhancers – *cis*-regulatory sequences composed of TF binding sites that, when bound, regulate expression of their target genes. Tens of immune regulatory sequences have been identified by searching for NF-κB and other motifs upstream of genes that encode antimicrobial peptides and other highly induced immune genes (Senger et al. 2004; Busse et al. 2007). However, the vast majority of enhancers that control the ∼1,000 immune-responsive genes are unknown. Identifying and investigating enhancers that regulate the innate immune system is critical to understanding how innate immunity coordinates a complex response upon infection. Flies provide an innate-only immune system to study these pathways, which are conserved not only across insects, but also in mammals (Dushay and Eldon 1998; Yu et al. 2022).

To identify immune regulated enhancers genome wide, we used STARR-seq (Self Transcribing Active Regulatory-Region sequencing) (Arnold et al. 2013; Muerdter et al. 2015). This high-throughput activity-based assay identifies sequences that act as enhancers to drive their own transcription. We performed this assay in S2* cells, a *D. melanogaster* hemocyte-like cell line that is more immune responsive than its parental line, S2 cells (Samakovlis et al. 1992; Cherbas et al. 2011). We focused on the immune response to a gram-negative bacterium, as it induces an immune response in a cell autonomous manner that can be reproduced in cell culture. To induce this immune response in S2* cells, we pretreated cells with the steroid hormone 20- Hydroxyecdysone (20E), before addition of heat-killed *Serratia marcescens* (HKSM), a gram-negative bacterium. In whole organisms, 20E has been shown to regulate development as well as immunity, and treatment of S2* cells with 20E induces expression of PGRP-LC, a critical receptor for DAP-type peptidoglycan in the Imd pathway (Rus et al. 2013).

Here, we report thousands of novel immune-responsive enhancers and dissect their TF composition and activity upon immune stimulation. We find that TF binding site content varies based on the function and timing of enhancer target genes. Relish, in particular, has a distinct role in enhancers active early in an infection and is enriched in enhancers regulating effectors. Effector genes are controlled by more immune specific enhancers, while signaling components are enriched for constitutively active enhancers. ATAC-seq in hemocytes upon immune stimulation reveals that while most enhancers do not change accessibility, those that are opened are enriched for specific transcription factor binding motifs including the ecdysone responsive EcR/Usp complex. Global documentation of immune enhancers in *Drosophila* reveals novel immune enhancers, with differential TFBS composition chromatin accessibility states.

## Results

### A STARR-seq assay reveals thousands of immune-responsive enhancers

To identify the enhancers that regulate the *Drosophila* innate immune response, we performed a STARR-seq experiment in S2* cells, a *Drosophila melanogaster* hemocyte-like cell line that is immune responsive (Samakovlis et al. 1992; Cherbas et al. 2011). Briefly, a genome-wide STARR-seq library of plasmids containing 500-750 bp fragments, under control of the *Drosophila* synthetic core promoter (DSCP), was introduced to cells via electroporation. After electroporation, cells were either treated with water, or the hormone 20-ecdysone (20E) to prepare them for immune stimulation. 24 hours after 20E treatment cells were treated with either PBS or heat-killed *S. marcescens* (HKSM). After 24 hours of immune stimulation, cells were collected and RNA was extracted, creating three treatment conditions, Control (water and PBS), 20E (20E and PBS) and IMD (20E and HKSM) (Fig. 1A).

**Figure 1:**
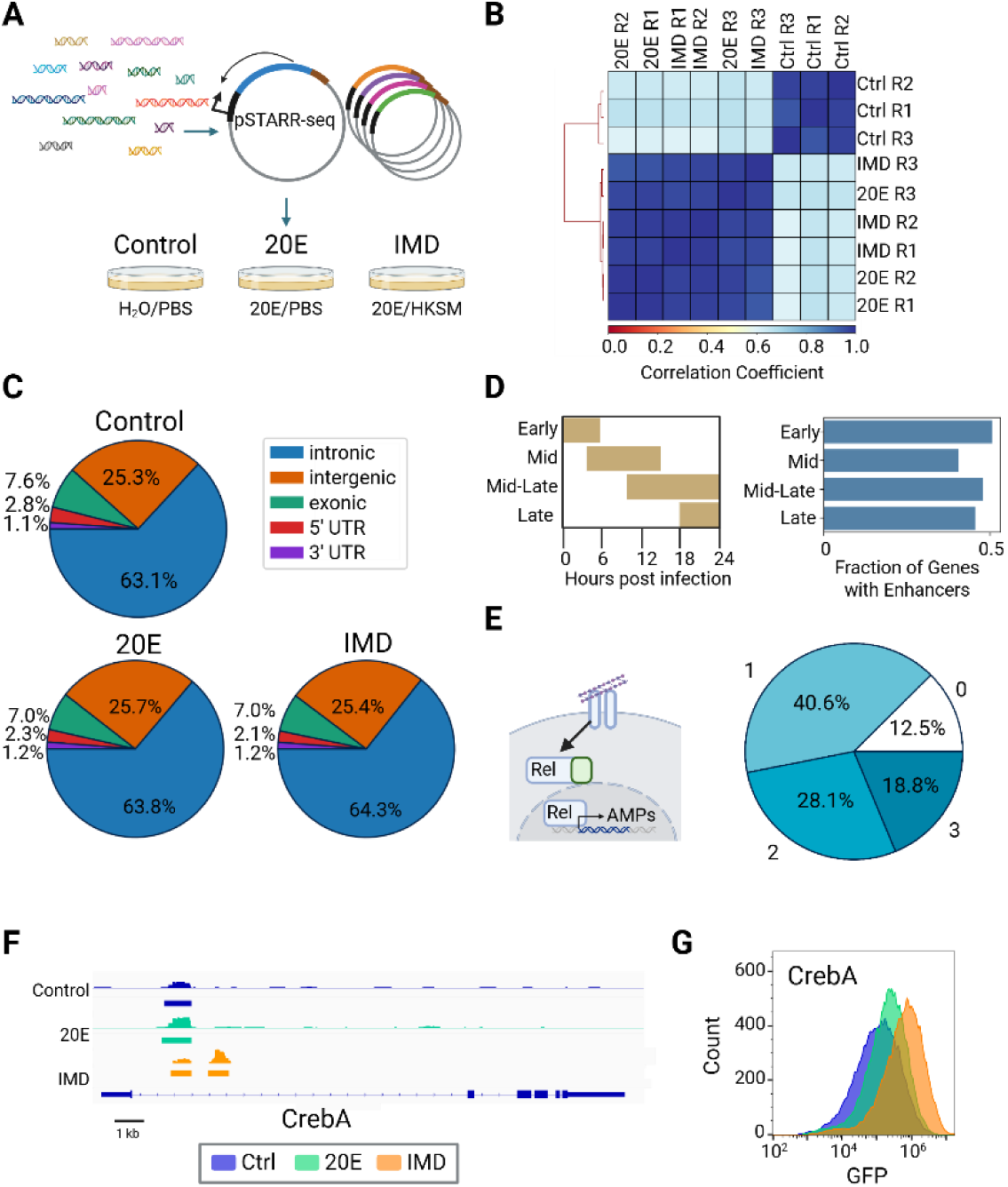
Immune enhancers identified by STARR-seq (A) Schematic of STARR-seq experiment. Fragmented genomic DNA is cloned into the pSTARR-seq plasmid, creating a library of putative enhancer regions. This library is transfected into S2* cells, split into 3 different populations and treated with either H2O/PBS, 20E/PBS, 20E/Heat-killed *Serratia marcescens* (HKSM). (B) Pearson correlation of each individual replicate across three treatments. (C) Distribution of enhancers in genomic regions (D) Schlamp et al. (2021) identified four time clusters of gene expression in the first 24 hrs following immune stimulation in adult flies; peak expression times for each time cluster are shown. Enhancers were assigned to these immune genes and the fraction of immune genes identified by Schlamp et al. with an immune enhancer is plotted. (E) Schematic of Imd pathway activated by DAP-type peptidoglycan binding to peptidoglycan recognition proteins (PGRPs) leading to the cleavage of the NF-κB Relish, allowing its translocation into the nucleus where it activates transcription of antimicrobial peptides (AMPs). Pie chart of the number of enhancers assigned to Imd-associated AMPs and effectors (Westlake et al. 2024) (F) IGV view of the CrebA gene locus, showing the STARR-seq output tracks and STARRPeaker called peaks for all treatment conditions. STARR-seq identified two enhancers in the CrebA intron, one shared across all conditions and one that is IMD specific (G) Overlay histogram of GFP levels of S2* cells transfected with a GFP reporter of the IMD-specific CrebA enhancer analyzed by flow cytometry.

Samples were processed following the STARR-seq protocol and sequenced along with the input plasmid library. Peaks of enhancer activity were called with the STARRPeaker program (Lee et al. 2020). Peaks from different replicates of the same treatment show greater than 0.95 Pearson correlation between them. Samples treated in the 20E and IMD treatment groups also show a high level of correlation, distinguishing them from Control treated samples (Fig. 1B).

To devise a set of enhancers that agreed across the replicates, we created the consensus enhancers, which contain the union of enhancers that have sequence overlaps at least 2 of the 3 replicates (Supplementary Fig. S1A). The consensus enhancer set contains 2,388 Control enhancers, 3,080 20E enhancers, and 2,934 IMD enhancers. Enhancers have an average length of 725, 746, and 743 bps for Control, 20E and IMD data sets, respectively. The shortest enhancers called by STARRPeaker are 500 bp, and the length distributions have a long tail, with the longest enhancers reaching over 2000 bps in length (Supplementary Fig. S1B). The consensus enhancers will now be used to represent the enhancers for each data set.

We next compared our consensus enhancers to previously-described enhancers to validate the properties of the enhancers identified by this STARR-seq experiment. To determine the distributions of enhancers across the genome, we binned enhancers by their genomic region. The majority (63.1%-64.3%) of these enhancers fall within intronic regions of the genome, with a smaller group (25.3-25.7%) found in intergenic regions, similar distributions to *Drosophila* enhancers found previously through STARR-seq (Arnold et al. 2013) or regions with classic enhancers chromatin marks, H3K4me1 and H3K27ac (Consortium et al. 2011) (Fig. 1C). Additionally, since S2* are derived from S2 cells, we asked whether the enhancers in the unperturbed sample (Control) aligned with enhancers identified in unperturbed S2 cells. 72% (1,727/2,388) of Control enhancers from the S2* cells have an overlap of at least 100 bp with STARR-seq enhancers found in S2 cells (Arnold et al. 2013). Overall, the consensus enhancers resemble previously identified *Drosophila* enhancers, in both sequence and genomic regions.

To determine if we identified enhancers that regulated immune genes, we assigned enhancers to genes. Since the vast majority of enhancers in *Drosophila* appear close to the genes they regulate, for each gene we assigned enhancers that occurred in the sequence spanning from 15 kb upstream to 5 kb downstream of the gene body, similar to other gene-enhancer assignments done in the *Drosophila* genome (Arnold et al. 2013; Kvon et al. 2014). Previous work has finely mapped the time-course of transcriptional upregulation of immune genes in the first 24 hrs of Imd stimulation in adult flies (Schlamp et al. 2021). These genes were then clustered by their expression interval. After assigning enhancers to these genes, we found IMD enhancers for just under half of these genes (252/551), fairly evenly distributed across the time clusters (Fig. 1D). Given that this list includes genes involved in other branches of the immune response, including the Toll and antiviral signaling pathways, we did not expect to find enhancers for the complete list of genes. When looking at a core list of Imd-associated effector peptides, including antimicrobial peptides, we found IMD enhancers near almost all of these genes (28/32) (Fig. 1E) (Westlake et al. 2024). Therefore, we conclude that enhancers found in the IMD set likely control known immune genes.

As an illustration of the data, we found two novel enhancers in the first intron of the transcription factor CrebA – one shared across conditions and one that is IMD-specific (Fig. 1F). CrebA is upregulated upon infection by both the Toll and Imd pathways and is crucial for survival after infection with a diverse set of bacteria because it prevents infection-associated ER stress (Troha et al. 2018). To confirm the IMD-specific enhancer’s activity and response to immune stimulus, we expressed GFP under the control of this enhancer and a minimal promoter and introduced this single reporter into S2* cells. We saw an increase in mean GFP expression by flow cytometry in IMD treated cells as compared to 20E and Control cells (Fig. 1G). The features of the enhancers identified by STARR-seq, including the close association of IMD enhancers with known immune genes and validation of an individual enhancer, suggest that our approach has identified a large set of *bona fide* enhancers.

### Analysis of enhancers reveals a variety of TFs used to activate each category of enhancers

To find the transcription factors (TFs) controlling immune enhancers, we searched for the motifs that are enriched in the IMD enhancer set. We used i-cisTarget, a web tool that searches through a database of over 130,000 regulatory features, including position weight matrices, to identify features overrepresented in a set of sequences, as compared to the genomic background (Imrichová et al. 2015). We first identified motifs enriched across the entire set of IMD enhancers, which yielded 473 significant motifs. Many of these motifs are similar, so we grouped them by sequence similarity using phylogenetic trees and examined the motif tree for TF families (Supplementary Fig. S3A). We selected TF families for further analysis based on several criteria. First, we selected the TFs with well-established roles in Imd immunity (Relish and the heterodimer pair Kay/Jra (Tafesh-Edwards and Eleftherianos 2020)) and ecdysone response (EcR/Usp and Eip74EF) (Rus et al. 2013). Next, we identified enriched motifs for TFs and TF families that have roles in hemocyte differentiation: GATA factors, forkhead TFs, and Gcm. GATA motifs can be bound by several family members, so to identify which TFs are likely active, we examined RNA-seq data generated alongside the STARR-seq experiment. *Srp* and *pnr* were both highly expressed in S2* cells and are required for hemocyte development (Spahn et al. 2014; Minakhina et al. 2011). Likewise, among the forkhead TFs, *jumu* is the most highly expressed and is required for hemocyte differentiation and phagocytosis in larvae (Hao et al. 2018). Gcm is a zinc finger transcription factor that is expressed in embryonic hemocytes and regulates their development (Bazzi et al. 2018).

To broaden our search for TFs, we used two approaches. First, we focused on motifs that were enriched in IMD enhancers i-cisTarget and whose associated TF is expressed in S2* cells, but without a reported immune role. These include Trl, the GAGA factor in *Drosophila*, as well as the bZIP TFs, Xbp1 and Xrp1. Second, we searched for motifs enriched in enhancers associated with genes in the four immune time clusters. We saw a number of basic helix-loop-helix (bHLH) motifs enriched in enhancers associated with the late time cluster (Supplementary Fig. S3B). We included SREBP because it is activated by gram-negative bacteria in the intestines and regulated by Imd in adipocytes, and Crp since it was highly enriched via i-cisTarget. Both of these bHLH transcription factors are also highly expressed in S2* cells. The motif for Hnf4, a zinc-finger nuclear hormone receptor expressed in fat bodies and in S2* cells, was also included (Barry and Thummel 2016). While not comprehensive, with this set of 13 TF motifs, we found at least three binding sites in each of the IMD enhancers (Fig. 2A, C), indicating that this TF set has a role in the majority of the 2,934 IMD enhancers.

**Figure 2:**
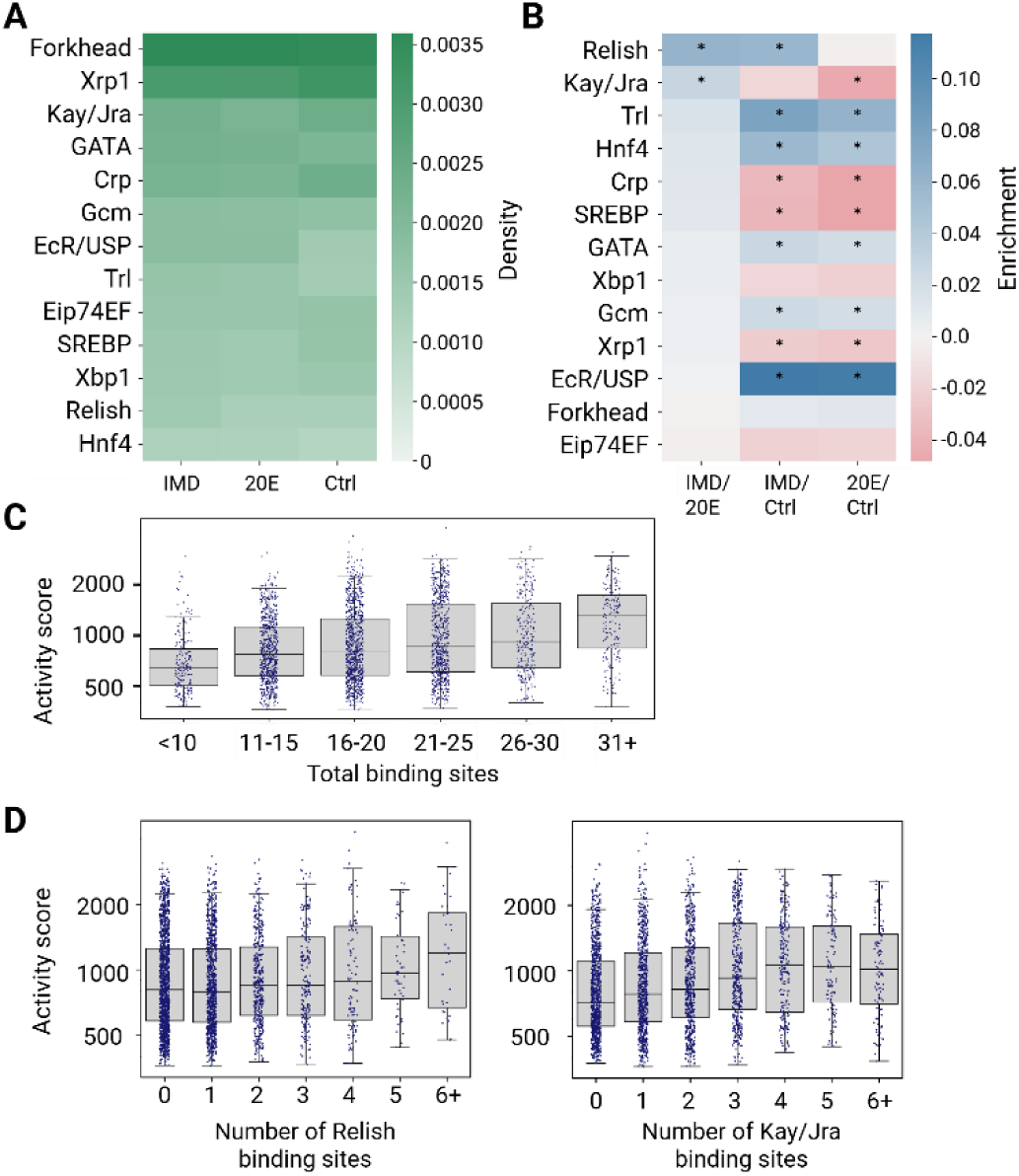
Transcription factor binding content varies between immune enhancers (A) Number of binding sites for each TF normalized by total length of enhancers in data set. (B) Log of odds ratio of TFBSs in three comparisons. Positive enrichment values are in blue and negative enrichment values are in red, and significance is indicated by * (p <0.05, Fisher’s exact test). (C) Activity score (log scale) from STARR-seq of IMD enhancers plotted by total binding sites. Individual enhancers are shown as individual circles and median and quartiles are displayed as box and whisker plots. (D) Activity score (log scale) of IMD enhancers by number of Relish or Kay/Jra binding sites.

Since our set of IMD enhancers also includes those that are constitutively active or induced upon ecdysone addition, to identify TFs with specific immune roles, we compared the density of TFBS between each treatment condition (Fig. 2B). When comparing 20E to the Control, we found the 20E enhancers are significantly enriched for EcR/Usp sites (p < 0.05; Fisher’s exact test) as expected, given the role of this complex in the ecdysone response. In the IMD condition, we expected to find enhancers that are activated from both individual responses to 20E and bacteria, as well as synergistic interactions between the two stimuli. Accordingly, we found a significant enrichment in motifs for immune TFs, i.e. Relish, Trl and Hnf4 in the IMD vs. Control comparison. To isolate the response solely to immunity (from treatment with HKSM), we then compared IMD enhancers to 20E enhancers and found significant enrichments in motifs for Relish and Kay/Jra, TFs known for their role downstream in the Imd pathway. Though the remainder of the TF motifs we analyzed are not significantly enriched in the IMD enhancers compared to the 20E enhancer, a subset is significantly depleted in the comparison of 20E enhancers versus the Control set. Some of these motifs may be more commonly found in constitutively active enhancers or enhancers that are active in the Control condition, but not the 20E or IMD conditions. Below, we also test the hypothesis that some of these TF play a role in specific subsets of immune enhancers with distinct functions (Fig. 3).

**Figure 3:**
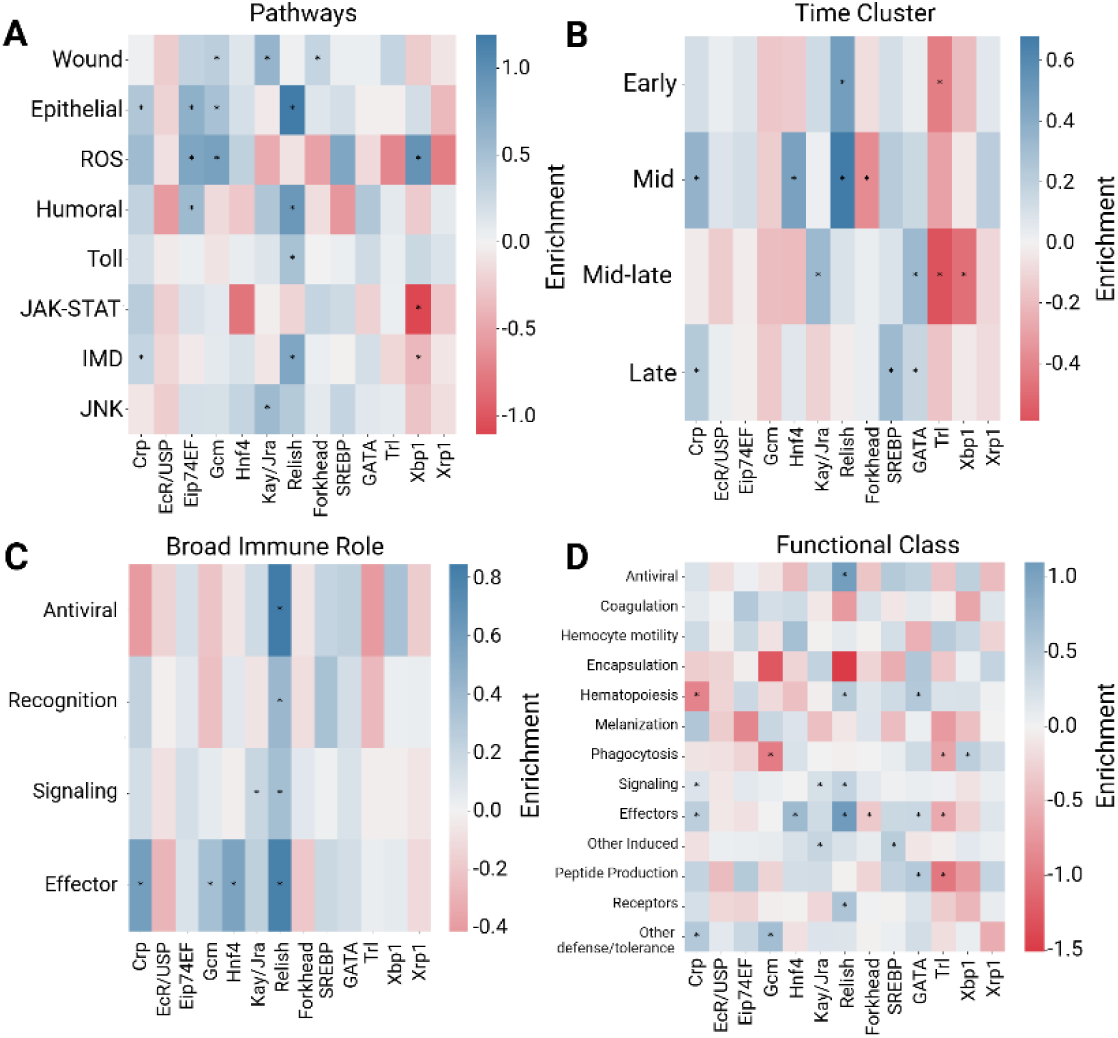
TFBS profiles of IMD enhancers vary according to pathways, timing, and function. Log odds ratio of TFBS in IMD enhancers grouped by pathway and process (Schlamp et al. 2021) (A) time cluster (Schlamp et al. 2021) (B), broad immune role (Schlamp et al. 2021) (C), or functional group (Westlake et al. 2024) (D) Positive enrichment values are in blue and negative enrichment values are in red, and significance is indicated by * (Fisher’s exact test).

We hypothesized that if these 13 TFs are functionally regulating the IMD enhancers, we would observe a correlation between the number of TFBS and the enhancer’s activity. The activity score estimates the strength of each enhancer as the fold-change between output RNA sequence and input DNA sequence in STARR-seq. We observed that as the number of TFBS per enhancer increases, activity score generally increases (Fig. 2C, Supplementary Fig. S2A, B) The top two enriched motifs, Relish and Kay/Jra, also show this relationship in IMD enhancers, with a general increase in activity score with more binding sites, although Kay/Jra plateaus around 4 TFBS (Fig. 2D). For the other 11 TFs in IMD enhancers, the relationship between activity score and number of binding sites varies; some TFs show a positive relationship, while others have no correlation (Supplementary Fig. S2C). Overall, we found a correlation between total binding sites and activity, indicating a functional role for these TF motifs in enhancer activity.

### Enhancers TF binding site content varies based on the function of the genes they regulate

After identifying TFBS in the IMD enhancer set, we asked whether the compositions of enhancers differ depending on the genes they regulate. We hypothesized that different sets of enhancers may be activated by distinct sets of TFs, depending on each set’s function, pathway, or timing of expression. We identified potential gene targets of each enhancer in an enhancer-centric fashion – by assigning all genes in the region ranging from 15kb upstream to 15kb downstream to that enhancer (Arnold et al. 2013; Kvon et al. 2014).

First, we grouped enhancers by immune pathways based on their assigned genes (Schlamp et al. 2021). As expected, we saw Relish sites enriched in Imd pathway related enhancers, as well as Kay/Jra enrichment in enhancers associated with the JNK pathway and wound response. ROS associated enhancers are enriched for Eip74EF, Gcm and Xbp1 motifs. Wound response enhancers are enriched for hemocyte related TF binding sites: Gcm and Forkhead TFs. Interestingly, we saw Relish also enriched in Toll pathway related enhancers suggesting some interactions between the Toll and Imd pathways (Fig. 3A). Transcription factor binding sites, e.g. Relish and Kay/Jra, are enriched in their corresponding pathways, demonstrating the utility of analyzing the general immune enhancer data set in smaller sets.

We next hypothesized that the enhancers that regulate genes induced at different times may rely on different TFBS. Since transcripts produced from the STARR-seq plasmid are stabilized via a poly-A tail, we expected to collect transcripts from enhancers that were active at any point within the first 24 hrs of immune induction. Using the finely mapped time-course data from the first 24 hrs of immune stimulation by Schlamp *et al*. 2021, we labeled enhancers with different time clusters based on the expression of their assigned genes. We found that Relish sites are enriched in enhancers that regulate genes expressed early to mid immune induction (0-16 hrs) when compared to all IMD enhancers (Fig. 3B). Late expressing genes, 12-24 hrs, are enriched for bHLH sites, SREBP and Crp, as well as GATA sites. SREBP is an activator of lipogenesis, typically triggered by low lipid levels, including during infection (Charroux and Royet 2022). Since Imd activation depletes lipid stores, its late activation may be in response to a metabolic shift. Nothing is yet reported on Crp’s potential role in the immune response – it is possible that this motif is representative of other bHLH TFs (including SREBP). However, *crp* is highly expressed in S2* cells. Therefore, this analysis reveals that there is distinct regulatory logic at different stages of the immune response, with early-acting enhancers regulated by the canonical Imd-responsive TF, Relish and later acting enhancers relying on bHLH activators, suggesting a mechanism for how genes are temporally regulated.

To determine if the role of a gene, regardless of pathway, affects how it is regulated, we investigated the composition of TFBS using two sets of gene annotations. Using broad immune role annotations, e.g. antiviral response, recognition, signaling, or effector, (Schlamp et al. 2021), we found effectors are controlled significantly by several TFs: Crp, Gcm, Hnf4, and Relish. The signaling and recognition stages of the immune response are also significantly controlled by Relish (Fig. 3C). We also used more detailed functional annotations from a diverse and comprehensive set of genes involved in the response to infection, categorized by their biological role in immunity (Westlake et al. 2024). We saw several TFs enriched within functional groups. We found that Relish is the most highly enriched transcription factor in enhancers assigned to effectors, followed by Hnf4 (Fig. 3D). Relish and GATA motifs are enriched among enhancers associated with hematopoiesis. Interestingly, Xbp1, a known component to the unfolded protein response (UPR) (Huang et al. 2017), is enriched in both reactive oxygen species (ROS) and phagocytosis related enhancers. Overall, we find that our set of immune-induced enhancers encompasses many facets of the immune response. Enhancer composition reflects differences in the timing, pathway and function of their target genes.

### Enhancers can be further divided into activity classes

Up to this point, we have analyzed all the enhancers active in the IMD condition together. However, it is possible that some of these enhancers are active in the Control and/or 20E conditions, while others are only active upon IMD stimulation. To discern between these types of enhancers, we grouped enhancers based on the specificity of their activity across all three conditions. Seven *activity classes* were determined: Constitutive, Control Only, 20E Only, IMD Only, Control + 20E, Control + IMD, and IMD + 20E. We found that the Constitutive enhancers comprised the largest activity class with 1,344 enhancers; the second largest group was IMD + 20E enhancers with 1,106 enhancers. Many enhancers are likely active in both the 20E and IMD conditions because both treatments include 20E, but we also found 372 enhancers that are active only in the IMD condition (Fig. 4A). We found relatively few enhancers active in the Control + IMD and Control + 20E groups, which is biologically sensible – it is hard to posit a mechanism that activates enhancers in the Control and IMD, but not 20E condition, or in the Control and 20E, but not IMD condition, due to the shared 20E stimulus in the IMD and 20E conditions.

**Figure 4:**
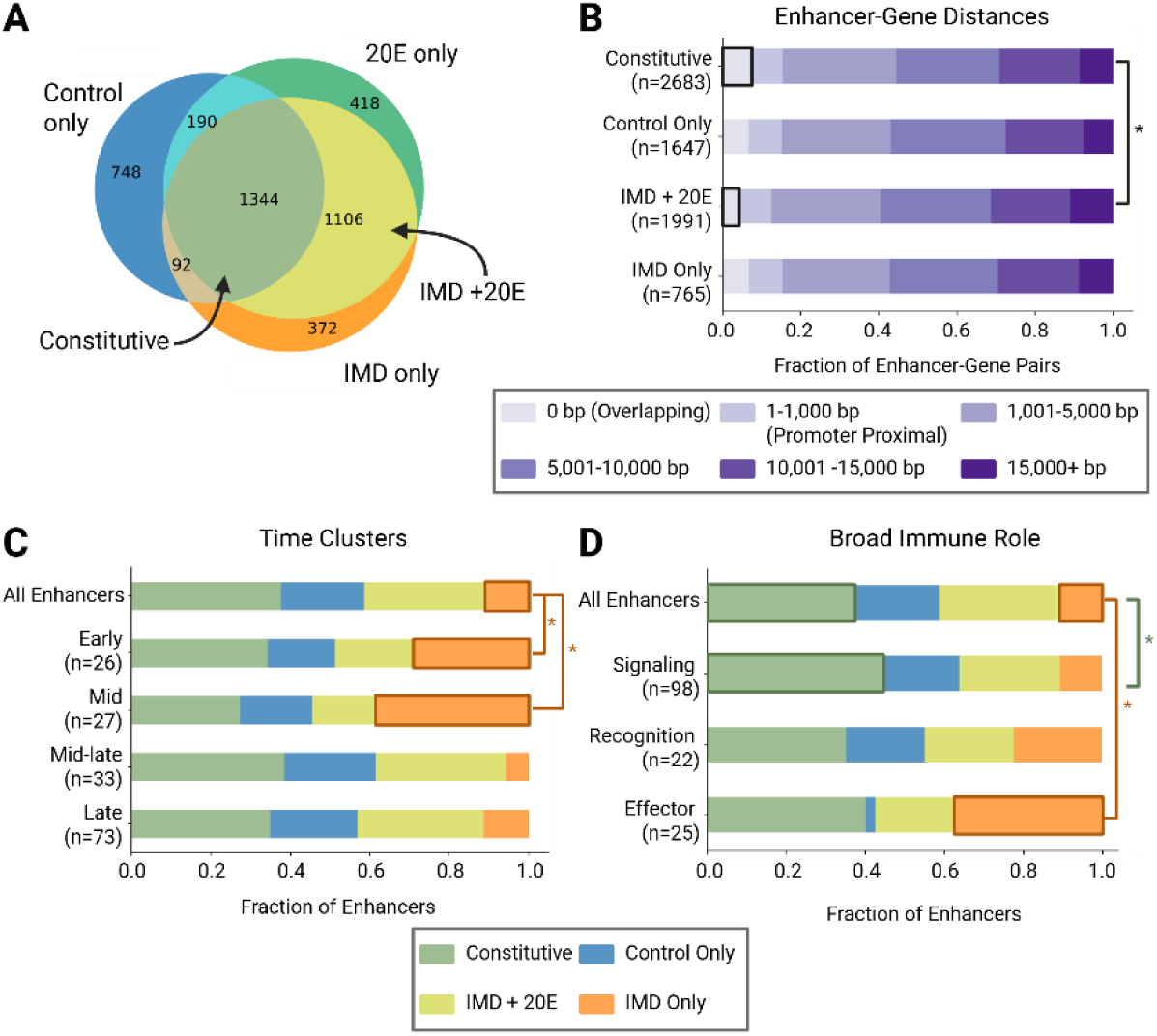
Activity classification of enhancers reveals specific properties of inducible and constitutive enhancers. (A) Venn diagram of enhancers showing activity in treatment conditions, defining activity classes. (B) Distributions of distances between enhancers and paired genes. Distance is measured from the closest edge of the enhancer to the transcriptional start site (TSS). The fraction of Constitutive enhancers that overlap with their paired genes is significantly greater than the fraction of inducible IMD + 20E enhancers that overlap with genes (2-sided proportion z-test, p-value < 0.05). The number of Enhancer-Gene pairs is listed in parentheses for each activity class. (C) Distribution of enhancer activity classes by time cluster (Schlamp et al 2021). The fraction of IMD Only enhancers is significantly larger in the Early and Mid clusters as compared to All Enhancers (one-proportion z-tests, p<0.05, indicated by *). The number of genes in each time cluster is listed in parentheses. (D) Distribution of enhancer activity classes by broad immune role (Schlamp et al. 2021). There is a significantly higher fraction of IMD Only enhancers in the effector group as compared to All Enhancers, while the signaling group has a higher fraction of constitutive enhancers as compared to All Enhancers (one-proportion z-tests, p<0.05, indicated by *).

To investigate the contributions of each TF in determining an enhancer’s activity class, we trained a logistic regression model to predict activity class based on the enhancers’ TFBS content. This model has a modest ability to discern between activity classes based on TF motif content, with a weighted average F1 score of 0.3 (maximum F1 score is 1; Supplementary Fig. S4A, C). When we trained a model only using enhancers from the Constitutive, Control Only, IMD Only, and IMD + 20E classes, the model performance improved, with a weighted average F1 score of 0.43 (Supplementary Fig. S4B, D). Although these models are not very accurate, we can examine the coefficients for each motif, and evaluate how it contributes to the determination of each class. For both versions of the model, we saw Relish, Trl, and Kay/Jra binding sites contribute the most to IMD Only enhancers. As expected, enhancers active in both IMD and 20E are controlled by the ecdysone response, with EcR/Usp binding sites having the highest coefficient for that class. Overall, TFBS assignments from 13 motifs are informative, but not sufficient for determining the activity class of these enhancers.

### Enhancers’ activity classes are associated with their target genes’ immune role

The assignment of enhancers to activity classes allowed us to make more refined gene-enhancer assignments by pairing these assignments with RNA-seq data taken alongside the STARR-seq data. For each activity class, we first identified genes expressed in that condition. For example, for the Constitutive class, we considered genes expressed in the Control, 20E, and IMD conditions, while for the IMD Only class, we considered genes expressed in the IMD condition, regardless of their expression in other conditions (see Methods and Supplementary Fig. S5A). We did not restrict, for example, IMD Only enhancers to genes only expressed in the IMD condition because it is possible that the gene is controlled by multiple enhancers with different activity, e.g. one IMD-inducible enhancer and a second constitutive enhancer (Fig. 1F). Using these expressed genes, we then made gene-enhancer assignments using the same 15 kb upstream/5 kb downstream windows as before.

To test for the accuracy of these assignments, we also identified genes that were differentially expressed either between the Control and 20E condition, the Control and IMD condition, or the IMD and 20E condition. Notably, this is distinct from our filtering process, which only requires a gene to be expressed in a condition, not necessarily differentially expressed between conditions. In each case, we found the majority of gene-enhancer assignments for these differentially expressed genes were to enhancers in the appropriate activity class (Supplementary Fig. S5D). For example, 77% of genes that show a greater than two-fold change in expression between the 20E and IMD conditions have at least one IMD Only enhancer. We also compared gene-enhancer assignments with and without the expressed-based filtering and found that the filtering decreased the number of likely spurious gene-enhancers assignments (Supplementary Fig. S5B, C). Therefore, we have confidence that filtering for genes based on expression yields a high-quality set of gene-enhancer assignments.

Using these expression-based gene-enhancer assignments, we hypothesized that genes with different roles in the immune system might be controlled by enhancers with different activity classes. We first compared the enhancers controlling immune recognition, signaling, and effector genes (Fig. 4D). We found that effector genes had the largest proportion (38%) of IMD Only enhancers, significantly more than the proportion of IMD Only enhancers among all enhancers (p<0.05, one proportion z-test). Signaling genes had the largest proportion of Constitutive enhancers (44%), likely because these components have more consistent expression across conditions, (p<0.05, one proportion two-sample z-test). We also compared the enhancer activity classes for genes sorted by their time of induction and found that genes induced in the Early and Mid time clusters have significantly more IMD Only enhancers than genes induced later in the infection (Fig. 4C). Although the later time clusters include genes that are immune induced, they may be controlled by a larger proportion of Constitutive enhancers for several reasons. For example, some genes may be activated by secondary signals, e.g. changes in metabolism, that have roles in other pathways. Other genes may be expressed in tissues not well-modeled by the hemocyte-like S2* cells. This analysis indicates that, depending on their immune role or expression timing, specific gene groups are associated with different enhancer activity classes.

This analysis also allows us to investigate the distribution of enhancer-gene distances. We hypothesized that Constitutive enhancers were more likely to overlap with the transcription start site (TSS) of their target genes, since distal enhancers are often found to be condition- or tissue-specific (Zabidi et al. 2015). To address this, we measured the distance from the TSS of the gene to the closest edge of the enhancer, and binned these enhancer-gene pairs into six distance groups (Fig. 4B). Consistent with our hypothesis, Constitutive enhancers overlapped with the TSS of their target genes (the 0 bp group) more often than the inducible IMD + 20E enhancers (p<0.05, 2-sided z-test). But overall, we saw little difference in the full distribution of gene-enhancer distances based on activity class, with more than 60% of enhancers within 10,000 bps of the TSS of their target gene and 90% of enhancers within 15,000 bps.

### Individual enhancer reporters validate activity classes and suggest tradeoffs between IMD and Toll induction

To further validate the STARR-seq enhancer measurements and our assignments of enhancers to activity classes, we tested the activity of eleven enhancer reporters that place GFP under the control of the minimal *Drosophila* synthetic core promoter (DSCP) and an enhancer from the STARR-seq experiment (Fig. 5E). For simplicity, enhancers are named for the gene with the closest TSS that is expressed in the same conditions as the enhancer (with the exception of *dso* enhancer, see below). We chose nine immune-responsive enhancers from two groups: enhancers that aligned with previously identified immune enhancers and novel enhancers that were in the IMD Only or IMD + 20E enhancer classes. We also included one enhancer (*CG9837*) in the Constitutive class and the DSCP-only reporter as a negative control. We introduced the reporters into S2* cells individually and measured each reporter’s GFP expression levels in Control, 20E, and IMD conditions using flow cytometry (Fig. 5A). As expected, the reporters for the *PGRP-SD* and *Mtk* enhancers, which closely match previously identified enhancers from Busse et al. and Senger et al., respectively, significantly upregulate GFP upon IMD stimulation, as compared to the Control condition. Among the remaining seven reporters in the IMD Only or IMD + 20E classes, six enhancers upregulate transcription upon IMD stimulation (*CrebA*, *lectin-26Cb*, *RpS26*, *Lmpt*, *CG5758*, *Cog8*), and one enhancer does not (*dso*). As expected, the constitutive enhancer assigned to *CG9837* did not show increased activity upon IMD stimulation. To explore enhancer activity further, we characterized absolute, instead of fold change, reporter activity in the Control condition. The *CG9837* enhancer reporter drives high activity in both the Control and IMD conditions (Fig. 5C), validating that it is a strong constitutive enhancer, and the negative control (DSCP) drove low expression in both conditions, indicating that it is a minimal, weak promoter.

**Figure 5:**
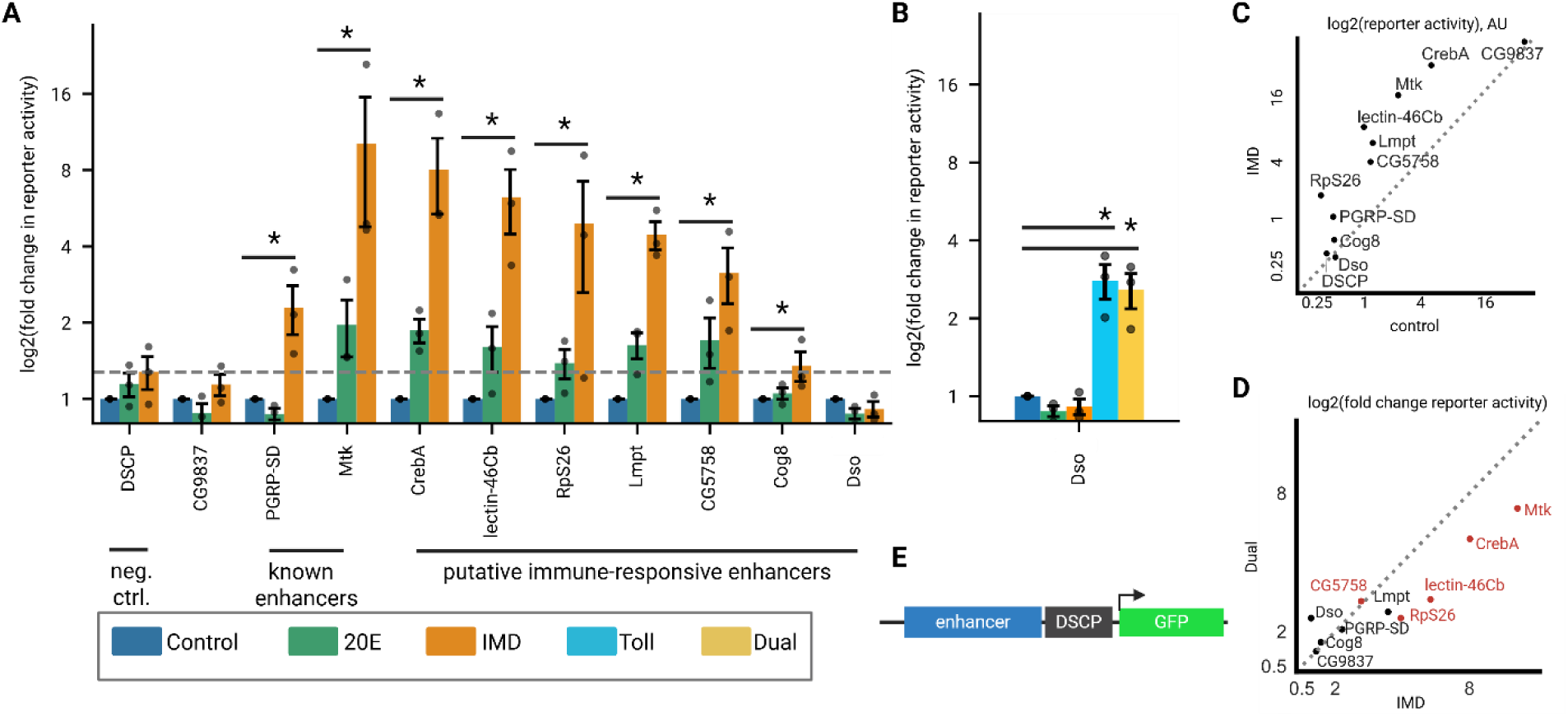
STARR-seq enhancers reporters demonstrate immune specific expression (A) Log_2_ (fold change) of GFP expression measured by flow cytometry normalized to Control treatment of enhancer reporters and promoter-only (DSCP) negative control. Bars represent the mean and dots show three biological replicates. Error bars represent standard error of the mean. Gray dashed line marks the expression level of DSCP control upon IMD stimulation. Significant gene induction is labeled with * p<0.05 (one sided Mann-Whitney test). (B) Log_2_ (fold change) of GFP expression normalized to Control treatment of *daisho* enhancer reporter. Significant gene induction is labeled with * p<0.05 (one sided Mann-Whitney test) (C) Log_2_ of GFP expression levels for reporters comparing IMD treatment to Control treatment. Both treatment conditions are normalized to untransfected S2* Torso-pelle cells. Dashed line is y=x, denoting an equal response to IMD and control treatment. (D) Log 2 of GFP expression levels for reporters comparing Dual treatment to IMD treatment. Both treatment conditions are normalized to Control treatment. Dashed line is y=x, denoting an equal response to IMD and Dual treatment. Enhancers highlighted in red are both Toll and IMD responsive (E) Schematic of enhancer reporter plasmids, with GFP under control of STARR-seq enhancers and minimal promoter, DSCP.

We were surprised to find an IMD-induced enhancer between *dso1* and *dso2* genes, since these genes are induced by the Toll pathway (Levy et al. 2004; Cohen et al. 2020). We chose to refer to this enhancer as *dso*, since it lies between *dso1* and *dso2* and the closest gene highly expressed in the IMD condition, Ipk1, does not have an immune function (Supplementary Fig. 6B). The *dso* enhancer activates GFP expression in response to Toll stimulus, either on its own or when both the IMD and Toll pathways are induced in the Dual condition (Fig. 5B), indicating this is a Toll-responsive enhancer likely controlling the *dso1/2* genes. Five of the eight IMD-responsive enhancers, *Mtk*, *CrebA*, *CG5758*, *lectin-46Cb* and *RpS26*, also respond to Toll induction beyond the negative control (Supplementary Fig. S6A), indicating that some enhancers can respond to both the IMD and Toll pathways.

Synthesizing the data across these experiments, we see several groups of enhancers: those that are only IMD-responsive (*PGRP-SD*, *Lmpt*, *Cog8*), those that are only Toll-responsive (*dso*), those that are Toll- and IMD-responsive (*Mtk*, *CrebA*, *CG5758*, *lectin-46Cb* and *RpS26*), and those that are constitutive (*CG9837*). To test if there is a tradeoff or synergy in enhancer function upon stimulation of the Imd and Toll pathways, we compare enhancer function in the IMD and Dual induction conditions (Fig. 4D). Among the Toll- and IMD-responsive enhancers (highlighted in red), we found that four of the five are more active in the IMD than Dual induction condition, indicating that the addition of a different stimulus generally has a cost to IMD-induced stimulation, possibly due to an increased burden on a cell’s resources. One enhancer, *CG5758*, is equally active in both Toll induction and IMD induction, avoiding this tradeoff. STARR-seq identified enhancers generally recapitulate their activity in individual enhancer-reporter constructs and demonstrate tradeoffs of inducing both the Imd and Toll pathways.

### Most enhancers do not change chromatin accessibility upon immune stimulation

The STARR-seq assay identifies sequences that can drive expression in the cell type and conditions tested, independent of chromatin context. In a whole organism, both this sequence information and chromatin structure determine an enhancer’s ability to regulate gene expression. To compare STARR-seq enhancers to the chromatin state in the animal itself, we conducted ATAC-seq in hemocytes isolated from adult flies in either control or immune-stimulated conditions. To mimic the cell culture induction and avoid contaminating bacterial DNA in the ATAC-seq library, we performed the immune induction using heat-killed *S. marcescens* (HKSM) and measured chromatin accessibility 24 hours after treatment (Fig. 6A).

**Figure 6:**
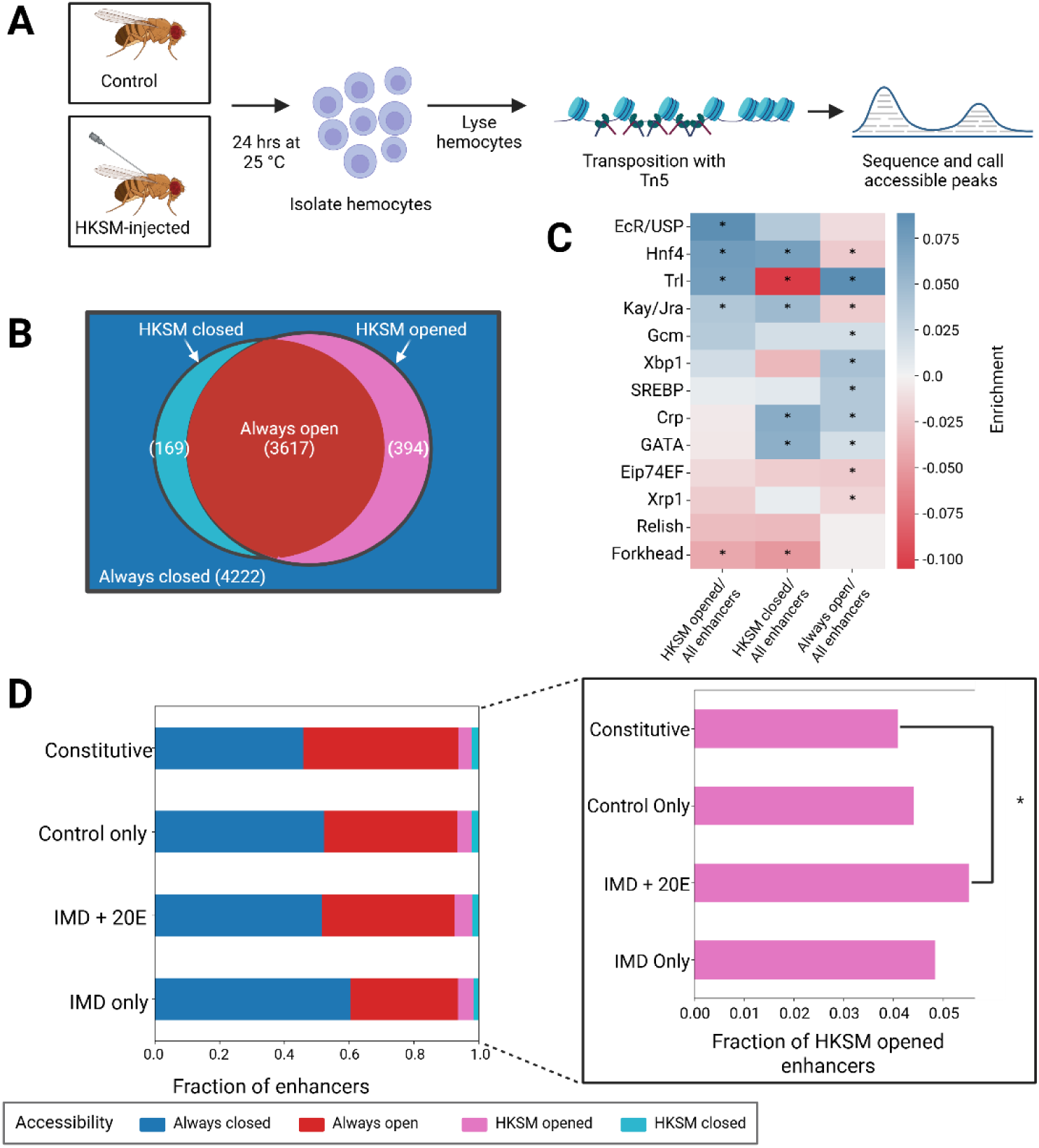
ATAC-seq reveals immune inducible enhancers are made accessible by EcR/USP, Hnf4 and Trl (A) Diagram of ATAC-seq on immune stimulated hemocytes. Flies are injected with HKSM and along with age matched controls are incubated at 25°C for 24 hours. Then flies are anesthetized, hemocytes are extracted via centrifugation before lysis and transposition. Tn5 was used to insert adaptors into accessible chromatin. These fragments are then isolated and sequenced to identify accessible peaks. (B) Venn Diagram of accessibility groups of enhancers. Always closed enhancers are in blue and are inaccessible in both the control and HKSM samples. The right circle is enhancers open in the control sample and the left circle is enhancers open in the HKSM sample. The intersection has enhancers open in both. The number of enhancers in each group is listed in parentheses. (C) Log odds ratio of TFBS in enhancers by accessibility compared to all enhancers, for previously identified motifs Positive enrichment values are in blue and negative enrichment values are in red, and significance is indicated by * ( p <0.05, Fisher’s test). (D) Fraction of enhancers of each accessibility group for biologically relevant activity classes. Insert of the same data for HKSM opened enhancers, plotted separately for clarity. A greater proportion of IMD + 20E enhancers are HKSM opened than Constitutive enhancers (* indicates p < 0.1, Two sample z-test).

In the original STARR-seq paper, roughly half of the enhancers were found to be accessible in the genome of the same S2 cell line used for the assay (Arnold et al. 2013). Here, we compared STARR-seq enhancers to the equivalent cell type in intact animals, and therefore expected that the overlap between enhancers and open chromatin might be slightly lower. However, we found that half (49.8%) of all STARR-seq identified enhancers are in regions that were ever accessible in hemocytes, similar to the percentage of previous STARR-seq enhancers found to be accessible in S2 cells. We then divided enhancers into accessibility groups: those that are open in both control and stimulated conditions (always open), those closed in both conditions (always closed), opened by HKSM stimulus, or closed by HKSM stimulus. More than >90% of enhancers maintain their chromatin structure for the duration of the experiment, either remaining open or closed (Fig. 6B). In enhancers grouped by activity class, we found that Constitutive enhancers are more likely to be always open than in IMD Only and IMD + 20E enhancers (Fig. 6D; p < 0.002, z-test, Bonferroni multiple test correction). The fraction of enhancers opened by HKSM is marginally higher in the IMD + 20E enhancers than Constitutive enhancers (p = 0.099, z-test). Similarly, when enhancers are grouped by treatment, there is a slight increase in HKSM opened enhancers in 20E enhancers vs. Control (p = 0.055, z-test) (Supplementary Fig. S7A). Therefore, the chromatin state of enhancers in the adult hemocyte is somewhat related with their STARR-seq activity, with a small fraction of immune-responsive enhancers lying in closed chromatin until an immune stimulus is present. However, the chromatin state of most enhancers identified via STARR-seq is unchanged in the 24 hours after treatment with HKSM.

To understand what determines the accessibility of different groups of enhancers, we measured the density of the 13 TF motifs we previously identified (Fig. 2) in each accessibility group. We found that compared to all enhancers, enhancers that are opened upon HKSM treatment are enriched for EcR/Usp, Hnf4, and Trl TFBS (Fig. 6C). Trl, a pioneer transcription factor known to open closed chromatin (Chetverina et al. 2021), is strongly enriched in HKSM opened and always open enhancers and strongly depleted in HKSM closed enhancers. EcR/Usp, which controls the ecdysone response, a key precursor to the Imd immune response, was found to be significantly enriched in the HKSM opened enhancers, but not in any other accessibility group. Previous work has identified Hnf4 as a key regulator of metabolism and lipid storage in development (Barry and Thummel 2016; Storelli et al. 2019). Here we saw its binding sites enriched in enhancers that either opened or closed upon immune stimulation, possibly indicating that Hnf4 is performing its metabolic switch role in reaction to infection. Relish is not enriched in any accessibility group, indicating that its role is primarily activating transcription without influencing chromatin state. Additionally, we looked for motifs enriched in the accessibility groups through i-cisTarget. Most of the identified motifs overlapped with the 13 motifs already analyzed, but motifs for CrebA, an immune TF (Troha et al. 2018), and a putative zinc finger TF were differentially enriched between the accessibility groups (Supplementary Fig. S7B). Overall, we see that TFBS content is correlated with accessibility of enhancers and that EcR/Usp may play a role in opening chromatin for enhancers in response to immune stimulation.

## Discussion

In this study, we have identified immune responsive enhancers across the entire *Drosophila* genome. Immune enhancers are enriched in motifs for Relish and Kay/Jra, the terminal transcription factors of the Imd and JNK pathways, respectively. We also find that distinct classes of immune enhancers differ by their TFBS composition and activity class. While most enhancers’ accessibility does not change upon infection in hemocytes, enhancers that are opened upon immune stimulation may be controlled by ecdysone signaling with an enrichment of EcR/Usp binding sites.

As was expected, Relish is the most enriched transcription factor in immune responsive enhancers, since the Imd pathway culminates in its activation. Here, we determine that Relish is a general activator of all Imd enhancers, but is preferentially found in the immune specific enhancers of early response genes and effectors. The Imd pathway activates the JNK pathway via Tak1 and concordantly, we find Kay/Jra binding sites enriched in immune enhancers and particularly in late acting enhancers (Tafesh-Edwards and Eleftherianos 2020).

The major pathways controlling the humoral immune response, Toll and Imd, have long been described as independent, each activated by different microbes, but there has been evidence of interactions between the pathways. For example, wounding and damage associated signals activate both immune pathways (Troha et al. 2018; Schlamp et al. 2021). Heterodimers between Relish and Dif, an NF-κB protein downstream of Toll, can form and activate transcription, incorporating signals from both pathways (Tanji et al. 2007, 2010; Morris et al. 2016). Also, certain genes like *Mtk*, *Drs*, and *CrebA* can be activated by both Toll and Imd (Busse et al. 2007; De Gregorio et al. 2002; Troha et al. 2018; Valanne et al. 2010). Consistent with this observation, we see that both Imd and Toll activate the enhancers for *Mtk* and *CrebA*. But, there is no evidence of synergistic enhancer activity when both stimuli are present (Supplementary Fig. S6A). Regulation by both pathways could be encoded in enhancers in multiple ways. The enhancers could contain distinct binding sites for Relish and Dif, common sites that can be bound by Relish or Dif, or sites that can be bound by Relish-Dif heterodimers. A previously-studied *Mtk* enhancer has Relish-specific sites and a common site that was experimentally determined to bind both Dif and Relish (Busse et al. 2007). We also identified one Dif-specific site in our expanded *Mtk* STARR-seq enhancer, suggesting the gene’s activation by both pathways may be mediated by both distinct and common sites. In addition to three Relish-specific sites, the *CrebA* enhancer contains a “GGGAATTCT” site, which closely resembles a common NF-κB site, and “GGGAACCACT” site, which has a 1 bp mismatch with the Relish-Dif heterodimer site (Tanji et al. 2007, 2010). Relish’s enrichment among the enhancers of Toll pathway components (Fig. 3A) suggests that other immune enhancers integrate signals from multiple pathways. Overall, we find evidence that immune enhancers are a point where the Toll and Imd pathways interact and combine signals.

Beyond the roles for Relish and Kay/Jra, we find the patterns of motif enrichment for Xbp1, Trl, Crp, SREBP, and Hnf4. Xbp1 is enriched in enhancers associated with ROS and phagocytosis. related enhancers. Upon infection, a biphasic ROS response stimulates hemocytes to differentiate (Myers et al, 2018). Perhaps, an increase in secreted peptides during an immune response activates the UPR and Xbp1 could in turn regulate the ROS and phagocytic pathways. Trl motifs were not enriched in enhancers associated with any gene classes based on immune function or immune expression timing (Fig. 3), but were enriched in both always open and HKSM opened enhancers in hemocytes (Fig. 6C). This suggests that Trl’s role in the immune response is remodeling the chromatin landscape, consistent with its role in development (Gaskill et al. 2021). Additional experiments and analysis may clarify if the Trl is only recruited to the HKSM opened enhancers upon stimulation, and if so, what precludes it from binding prior to stimulus. Crp, a known anti-apoptosis factor (Sopko et al. 2015; Atkins et al. 2016), could function to maintain the hemocyte population upon infection, with motifs enriched in mid- and late-expressed immune genes and among enhancers associated with effector and signaling genes. Both Hnf4 and SREBP have roles in regulating lipid biosynthesis and metabolism, and *SREBP* is downregulated in *Hnf4* mutants, suggesting a link between these genes (Vonolfen et al. 2024). There is evidence that these TFs are differentially regulated during infection or inflammation from studies that focus on different cell types (Charroux and Royet 2022). Given the large metabolic impact of the immune response (Bland 2023), further studies into the activity of these TFs in different tissues and stages of infection may reveal how expression is regulated after the immediate response driven by canonical immune-responsive TFs, like Relish and Kay/Jra. Hnf4 motifs are enriched in enhancers associated with mid-induced genes, and SREBP motifs are enriched among late genes. In addition, our TF assignments rely on the availability and quality of motifs for *Drosophila* transcription factors; other transcription factors could regulate immune enhancers that we are unable to identify.

Our data emphasizes the significant role of the hormone ecdysone in modulating the transcriptional immune response. Ecdysone is the master regulator of *Drosophila* developmental transitions, but it is also required for the Imd response in both cells and whole animals (Meister and Richards 1996; Rus et al. 2013). Ecdysone, which activates the EcR/Usp TF complex, induces the expression of the peptidoglycan receptor PGRP-LC, increasing the responsiveness of cells to peptidoglycan from gram-negative bacteria. If ecdysone increased Imd signaling by only acting on upstream components of the pathway and not on activating enhancers, we would expect to see that the IMD + 20E activity class enhancers contain EcR/Usp and not Relish motifs. Instead, we find that just under half (45.3%) of IMD + 20E enhancers contain both EcR/Usp and Relish motifs. The co-occurrence of Relish and EcR/Usp sites suggest that ecdysone also has a role in activating Relish-dependent immune enhancers, a separate function from its function inducing upstream pathway components. Additionally, chromatin accessibility changes in hemocytes from adult flies upon immune stimulus reveal that EcR/Usp is highly enriched in enhancers opened by HKSM. EcR/Usp recruits TRR and the MLR COMPASS complex, which monomethylates H3K4, priming these enhancers for an immune response (Zraly et al. 2021). Therefore, we find that ecdysone and EcR/Usp are crucial regulators of transcriptional immune response likely by both increasing enhancer activation and chromatin accessibility.

Our *in vivo* ATAC-seq experiments in hemocytes revealed that roughly 90% of enhancers do not change their accessibility status during the course of the experiment, while the remaining ∼10% change. This relatively stable chromatin profile is in line with the model that the majority of enhancers available to a cell type acquire their enhancer histone marks (e.g. H3K4me1) during lineage specification, while a subset, so-called “latent enhancers,” acquire these marks after a stimulus. In macrophages, ∼15% of enhancers are latent and only gain enhancer histone marks upon stimulus, e.g. with LPS (Ostuni et al. 2013). While latent enhancers were originally defined by their histone mark profiles, the same study found a strong correlation between the acquisition of enhancer histone marks and DNA accessibility, suggesting that our HKSM opened enhancers are *bona fide* latent enhancers (Fig. 6B).

Our analysis of the TF motifs enriched in the HKSM opened set revealed that a similar set of TFs (EcR/Usp, Hnf4, Trl, and Kay/Jra) bind to both HKSM opened enhancers and all enhancers active in the IMD condition (Fig. 2B), suggesting that latent enhancers largely rely on the same TFs as enhancers poised for activation. Notably, Relish motifs are not enriched in the HKSM opened enhancer set. Based on our analysis of enhancers associated with genes expressed in early, mid, and late time points immune stimulation, Relish is preferentially enriched in enhancers associated with early and mid time points (Fig. 3B). We therefore speculate that the lack of Relish enrichment in HKSM opened enhancers is because Relish enriched enhancers are likely already accessible as in hemocytes, where latent enhancers are associated with genes with slower activation times.

The diverse array of enhancers characterized in this study reflects the demands of an innate immune system. In animals without an adaptive immune system, the innate transcriptional response must encompass everything needed to defend against invading pathogens, while balancing a limited metabolic budget and maintaining housekeeping functions. Identification of immune enhancers across both the expanse of the genome and functional roles of the immune system adds to our understanding of how the immune response is controlled and the multiple types of enhancers involved.

## Methods

### Genome-wide reporter library construction

To generate a genome-wide reporter library, genomic DNA was isolated from both male and female *iso-1* flies. A solution of 100 ng/µL DNA was sonicated on ice with a Sonic Dismembrator sonicator (Thermo Fisher Scientific) using an 8-tip probe for 10 cycles of 15 sec on/59 sec off at 25% Amp. Fragments of size 500-750 bp were extracted from a gel. The genomic library preparation and STARR-seq protocol was done as described in Neumayr, *et al*. and outlined briefly below. The NEBNext End Prep kit (New England Biolabs) was used to process genomic fragments before PCR amplification with 8 cycles. These genomic fragments were then cloned into the pSTARR-seq fly plasmid (AddGene, #71499) using Gibson cloning.

The genomic library was transformed into MegaX DH10B *E. coli* (Thermo Fisher Scientific) by electroporation (exponential decay, 2kv, 25 μF, 200 ohms). The culture was grown overnight shaking at 37°C in LB with ampicillin. Cells were pelleted at 3,000xg for 30 mins. The plasmid library was extracted with the ZymoPURE II plasmid Maxi prep kit (Zymo Research).

### STARR-seq in S2* cells

To test the genomic library’s enhancer activity, we performed STARR-seq in S2* cells. S2* cells were a gift from Steven Wasserman and grown according to standard methods in complete Schneider’s media at 28°C. For three condition experiment, 2.8 * 10^9^ S2* cells were washed with electroporation buffer (MaxCyte) before resuspension in electroporation buffer to a total volume of 6 mL with input STARR-seq library (300 µg). Cells were electroporated in a R1000 processing assembly (MaxCyte) in the Maxcyte ATX using the S2 protocol. After electroporation, cells were added to 150 µL of basal serum-free Schneider’s media with a final concentration of 0.0256 U/µLof DNaseI in the center of a tissue culture dish. Cells were incubated with DNaseI for 30 mins at 28°C before complete Schneider’s media [Schneider’s *Drosophila* Medium (Fisher), 10% heat inactivated FBS (Fisher), 2 nM L-glutamine, 50U/mL Pen/Strep (Life Technologies.), 1% Fungizone (Life Technologies)] was added to stop the enzymatic activity.

Cells were split into three treatment groups: Control, 20E and IMD, treated with either water (Control) or a final concentration of 40 nM 20E (Sigma-Aldrich) (20E and IMD) for 24 hrs, and then PBS (Control and 20E) or heat-killed *S. marcescens* at a final concentration of 0.4 OD (IMD) for 24 hrs. Total RNA was isolated with RNAeasy prep kits (Qiagen) from which mRNA was purified using oligo-dT Dynabeads before digestion with DNaseI. Reverse transcription, followed by junction PCR and barcoding PCR were completed as described in Neumayr et al. 2019. Paired end Illumina sequencing was performed on each sample of 3 biological replicates and the input library. 75,000,000-122,000,000 reads were collected for each sample.

Alongside the STARR-seq replicates, RNA-seq samples were also created in triplicate. Untransfected S2* cells were treated in the same manner to create the same three treatment groups: Control, 20E and IMD. Total RNA was sent to Genewiz for polyA mRNA selection and library preparation before sequencing. At least 36,000,000 reads were collected for each sample.

### Computational analysis

#### Identifying STARR-seq peaks

STARR-seq reads were aligned to the dm6 genome using Bowtie and samtools. Deeptools was used to convert alignment files to bigWig coverage tracks. STARRPeaker was then used to compare the output STARR-seq files for each sample to the input DNA library (Lee et al. 2020). Based on a p-value threshold <0.05, STARRPeaker calls enhancer peaks for each replicate. It then ascribes an activity score (100* (output/normalized-input)) to each peak, where output is a measure of the number of reads in the enhancer peak, and normalized-input is the coverage of the enhancer peak in the input genomic reporter library.

With the STARRPeaker defined peaks, we built consensus peaks that overlapped between replicates of the same treatment group. If a peak was in at least two out of three replicates (with at least 1 bp of overlap), a consensus enhancer was defined using the union of any of the enhancers from individual replicates that occur at that location. Consensus enhancers were assigned the average of the activity scores of the replicate peaks that contribute to them.

Since many enhancers appear across multiple treatment conditions, we defined non-overlapping enhancers or activity classes by categorizing them into one of seven groups: Constitutive (Control, 20E, IMD), Control Only, 20E Only, IMD Only, IMD and 20E, 20E and Control, and IMD and Control. When comparing enhancers across treatment conditions, enhancers were considered active in two or more conditions if at least 60% of either enhancer was in the overlapping region. Coordinates for enhancers in overlapping groups were defined as the union of the treatment defined enhancers.

#### RNA-seq analysis

RNA-seq reads were aligned with bowtie2 and counts were generated with Subread’s featureCounts. Transcripts per million (TPM) were then calculated. To determine agreement between the triplicates and identify any outliers, principal component analysis (PCA) was completed. Sample 1, a control sample, was determined to be an outlier and was removed from the data set, leaving 2 control samples, 3 20E samples, and 3 IMD samples. Differential gene analysis was also completed by calculating the log fold change between the following conditions: IMD vs. 20E, IMD vs. Control, 20E vs. Control.

#### Genomic bins

To determine in which genomic regions the STARR-seq enhancers are, we compared the genomic coordinates of each enhancer to the annotated FlyBase FASTA files for exons, introns, 5’ UTR, 3’ UTR and intergenic regions. Since the *D. melanogaster* genome is very dense in genes, many enhancers occupied multiple genomic regions. To limit the region assignment to one region, we first chose the region with the highest overlap with the enhancer. If multiple regions have identical overlap lengths, we then choose the region based on the following hierarchy: 1. intergenic regions, 2. introns, 3. 5’ UTRs, 4. 3’ UTRs, 5. exons.

#### Transcription factor assignments

To identify transcription factor binding sites within all the IMD enhancers and within time-specific sets of IMD enhancers, we used i-cisTarget (version 6.0) to find motifs with a normalized enrichment score above 3.0 (Imrichová et al. 2015). Motifs were then grouped into motif trees using the R bioconductor package universalmotif. Motifs were selected based on several criteria: representation of a motif tree branch, coordinating transcription factors expression levels in S2* cells upon Imd stimulation, and known role in immunity or hemocytes. Once motifs were curated, they were identified in each Control, 20E and IMD enhancer with the FIMO tool of the MEME suite. FIMO scanned both strands of enhancers and TFBS with a p-value < 10^-3^ were chosen.

#### Gene assignments

Permissive gene assignments were made to enhancers by proximity. Gene centric enhancer assignments (used in Fig. 1) were made by scanning 15 kb upstream and 5 kb downstream of a gene and assigning any enhancer in this window to the gene. Enhancer centric assignments (as used in Fig. 3) scan 15 kb upstream to 15 kb downstream of an enhancer and assign any genes to that enhancer.

More restrictive enhancer-gene assignments were made by sorting genes into activity classes based on the fraction of RNA-seq samples where the genes were expressed (Transcripts Per Million (TPM) > 1). The criteria for the activity classes are as follows: Control Only (2/2 Control samples), 20E Only (2/3 20E samples), IMD Only (2/3 IMD samples), Control+20E (4/5 Control and 20E samples), Control+ IMD (4/5 Control and IMD samples) 20E+ IMD (5/6 20E and IMD samples), and Constitutive (7/8 all samples). Notably, we did not define any exclusion criteria (i.e., a gene expressed in every sample belongs to all of the activity classes and can be matched to any enhancer in any activity class).

For each activity class, genes were matched to the allowed set of enhancers by scanning 15 kb upstream and 5 kb downstream of the gene. Any enhancer that fell within this window and belonged to the set of allowable activity classes was matched to the gene. Matching was repeated for each of the enhancer activity classes, resulting in a list of enhancer-gene pairs categorized by enhancer’s activity class. Notably, since matching was done in a gene-centric manner, not every enhancer was matched to a gene; however, genes were allowed to have multiple enhancers and enhancers were allowed to regulate multiple genes.

Pathways, time, immune role and functional designations were designated to enhancers via their assigned genes. Enrichment of TFBS in each of these categories was determined by calculating the odds ratio of enhancers within the category having a particular motif compared to the whole IMD set. Statistical significance was determined with a Fisher’s exact test.

#### Logistic regression

Logistic regression models to determine the TFBS that contribute to activity classifications were made using scikit-learn library with 1000 maximum iterations, a test size of 0.2 and balanced class weights.

### Construction and flow cytometry of enhancer reporters

STARR-seq consensus enhancers were amplified from *iso-1* flies and cloned upstream of the Drosophila Synthetic Core Promoter (DSCP) (Pfeiffer et al. 2008; Arnold et al. 2013) driving EGFP into the section of the pAc5.1/V5-His-A (Invitrogen) plasmid containing the origin of replication and ampicillin resistance gene via Gibson cloning. See Supplementary Table 1 for regions cloned. Plasmids were designed using Benchling (www.benchling.com). TSS’s contained within cloned enhancers were mutated by site-directed mutagenesis. An additional plasmid, pKM7 (p-IEX-mCherry), (Russell et al. 2021) was used as a positive transfection control. Plasmids were prepared with the ZymoPURE II plasmid Midiprep kit (Zymo Research) and concentrated via ethanol precipitation.

Plasmids were electroporated using the Maxcyte ATX machine following the S2 cell protocol. To allow for either Toll or Imd stimulation, S2* cells stably transfected with pMT-Torso-pelle (Sun et al. 2002) were used for this experiment. Pelle, a key component of Toll signal transduction, is fused to the transmembrane domain of Torso and placed under control of the metallothionein promoter which is activated upon the addition of CuSO_4_. Ten million cells per sample were washed in 2 mL of electroporation buffer (MaxCyte), then concentrated into 100 μL of electroporation buffer. After washing and concentrating, cells were mixed with 5 μg of enhancer reporter DNA, and 5 μg of the mCherry transfection control and loaded into a well of a OC-100x2 processing assembly (MaxCyte). Directly following electroporation, transfected cells incubated at 28°C for 30 minutes in a droplet of 50 μL of basal Schneider’s *Drosophila* Medium and 0.5 μLof 1000 U/mL DNaseI. 5 mL of Schneider’s Complete Media were added to the cells in the T25 cell culture plate post incubation. Cells were then split across into five wells of a 6-well plate, for each of five treatment conditions: Control, 20E, IMD-induced, Toll-induced, or Toll- and IMD-(dual) induced.

Cells were then treated with either water (Control) or 40 nM 20E for 24 hrs (20E, IMD-, Toll- and dual-induced. Cells were treated with either 100 µL of PBS (Control and 20E conditions), 100 µL of OD=10 Heat-killed *Serritia marcescen*s (HKSM) to induce IMD, or a final concentration of 500 µM CuSO_4_ to induce Toll, or both HKSM and CuSO_4_ for dual induction.

Following a 24-hour induction period, cells were centrifuged for 5 minutes at 100xg to pellet the cells. Each pellet was resuspended in 1 mL of FACS buffer (1x Ca/Mg++ free PBS, 1mM EDTA, 25mM HEPES pH 7.0, 1% FBS). After sample preparation, cells were analyzed using either a FACSCalibur or CytoFLEX flow cytometer. Fluorescence signals from GFP and mCherry reporter constructs were collected using appropriate filter channels. Instrument settings and gating thresholds were established using non-fluorescent and untransfected control samples. After gating for live, transfected cells, the geometric mean of GFP signal in the treatment conditions (20E, IMD, Toll and Dual) was normalized to the geometric mean of GFP signal in the Control for each transfected plasmid, which is the fold change in reporter activity. To estimate the absolute reporter activity across different cytometers, we normalized the geometric mean for GFP signal in each condition to the geometric mean of GFP signal for untransfected cells. At least 10,000 cells were collected for each sample. Each plasmid was tested across three biological replicates.

### ATAC-seq with adult fly hemocytes

We injected 60-70 2-7 day old male OregonR flies with ∼50 nL of heat-killed *Serratia marcescens* at OD=0.5 in PBS and incubated them at 25°C for 24 hrs, alongside uninjected age matched controls. Hemolymph including hemocytes were collected by placing 20-25 anesthetized flies in 0.5 mL tubes with 3 20G needle holes at the bottom. Flies were spun at 6,000xg at 4°C for 1 min into ATAC-seq lysis buffer (Grandi et al. 2022).

Tagmentation and library prep protocols were adapted from Grandi et al. 2022, as follows. After centrifugation, hemolymph was combined within the treatment samples (injected and uninjected) respectively and mixed by pipetting 5x, followed by 3 min incubation on ice.

Samples were washed with 1 mL of wash buffer before centrifugation at 500xg for 10 mins at 4°C. Supernatant was removed and ATAC-seq Tn5 mix (1x TD buffer, 0.33x PBS, 0.01% digitonin, 0.1% Tween-20, 0.05 µL of Tn5/µL of mix) was added to the pellet. Samples were placed in a 37°C shaking incubator for 30 mins, with brief vortexing half way through. The Tn5 reaction was stopped with the addition of DNA binding buffer from the Monarch PCR clean up kit (New England Biolabs) and samples were frozen before the rest of the cleanup kit was completed. Barcoding PCR was completed with Illumina primers and qPCR quantification with NEBNext Library Quant Kit (New England Biolabs). Paired end Illumina sequencing was completed on three biological replicates per condition and at least 100 million reads were collected per sample.

### ATAC-seq analysis

ATAC-seq reads were collected per sample and trimmed using CutAdapt. Reads were then aligned to the genome using bowtie2. Data was filtered for duplicate reads using the Picard MarkDuplicates tool. Reads were adjusted for Tn5 offset using bamtools, and peaks were called using MACS2, using narrow peak calling and a p-value of 0.01. FrIP scores were calculated using bedtools intersect. Consensus ATAC-seq peaks were built from the widest margins of open regions present in at least 2 replicates. STARR-seq enhancers were then labeled open if greater than 50 bps of enhancer were in consensus ATAC-seq peaks.

## Data Access

All raw and processed sequencing data generated in this study have been submitted to the NCBI Gene Expression Omnibus (GEO; https://www.ncbi.nlm.nih.gov/geo/) under accession number GSE308695.

## Acknowledgements

We thank Benedetta D’Elia for help in developing the ATAC-seq protocol and Ila Rosen for help in preliminary plasmid construction. We thank Steven Wasserman for the S2* cells. This work was funded by NSF Award 2223888 (to ZW) TH was supported by NIH Award T32GM150533 and CS was supported by Boston University’s Undergraduate Research Opportunities Program and the Boston University STEM Pathways Program supported by Department of Defense DoD STEM FY20 Award HQ00342110008. The content is solely the responsibility of the authors and does not necessarily represent the official views of any of the funders. Figures created with BioRender.com

## Author Contributions

L.B.C. and Z.W. conceived the project and wrote the paper. L.B.C. performed the experiments and analyzed the data except for the following: T.H. developed activity classes and expression filtered gene assignment. C.S. created and tested the enhancer reporters. J.R.G. analyzed ATAC-seq data. L.B.C. and Z.W. supervised all experiments and analysis. Z.W. obtained funding for the project. All authors have read, edited, and approved the final manuscript.

