## Supplementary Information for "A genome-wide survey reveals a diverse array of enhancers coordinate the *Drosophila* innate immune response"

### <u>Supplementary Data List</u>

**Supplementary Figure S1: STARR-seq enhancers have similar lengths across treatments**

**Supplementary Figure S2: Activity score varies by number of TFBS**

**Supplementary Figure S3: Several TF families are enriched in IMD enhancers.**

**Supplementary Figure S4: Logistic Regression analysis highlights the role of some TFs in predicting activity class.**

**Supplementary Figure S5: Expression filtering improves enhancer-gene assignments (A) Overview of gene-enhancer matching workflow.**

**Supplementary Figure S6: Enhancer reporters are differentially responsive to 20E, IMD, Toll and Dual Induction.**

**Supplementary Figure S7: Most enhancers maintain chromatin structure upon immune stimulation.**

**Supplementary Table 1: Enhancer reporters**

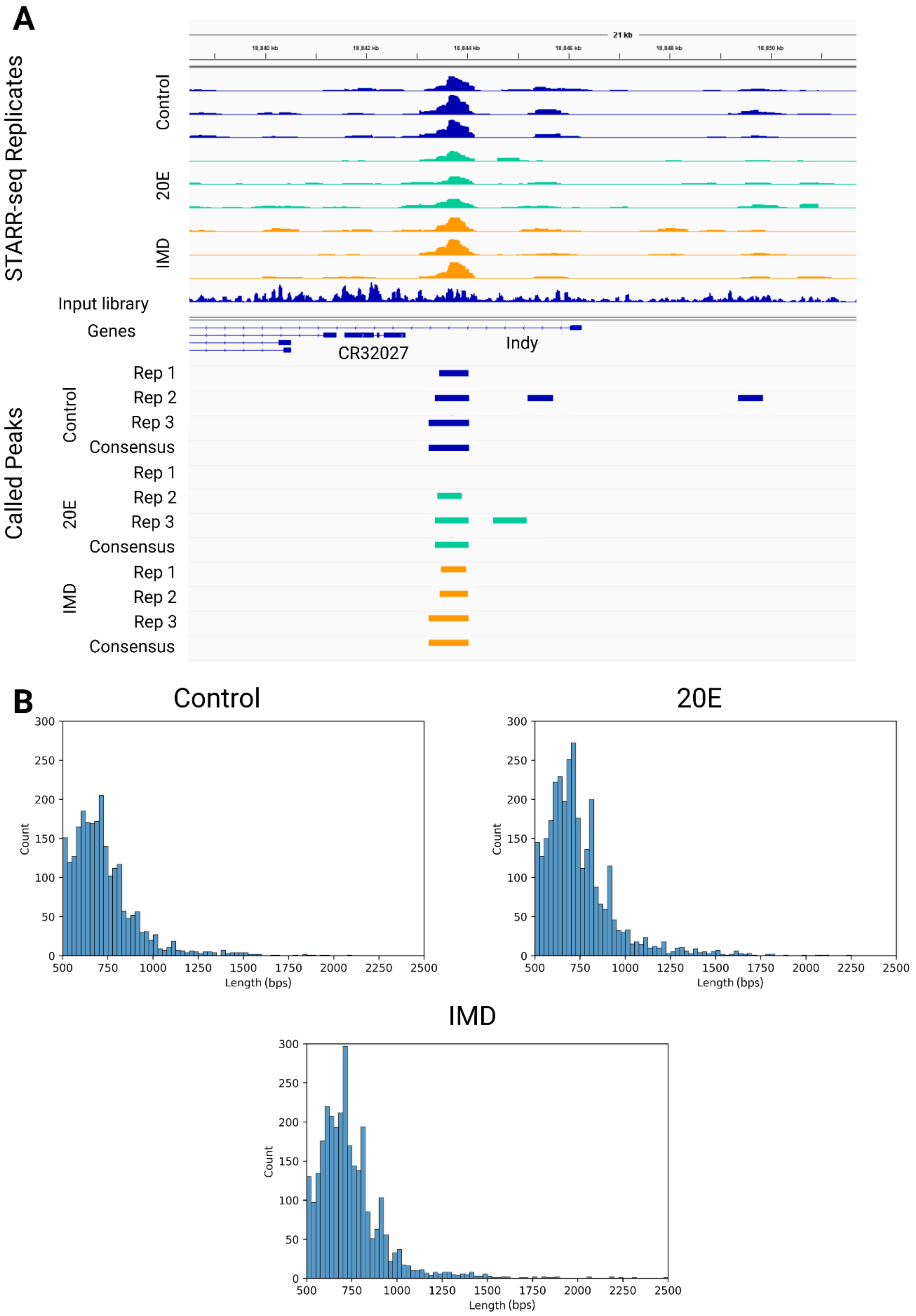


**Supplementary Figure S1: STARR-seq enhancers have similar lengths across treatments.** (A) IGV screenshot of tracks for STARR-seq replicates (output), input library, and references genes. STARRPeaker called peaks are shown for each replicate in addition to the consensus peaks for each condition. Control tracks are shown in blue, 20E tracks are shown in green and IMD peaks are shown in orange. (B) Histogram of lengths for STARR-seq consensus enhancers.

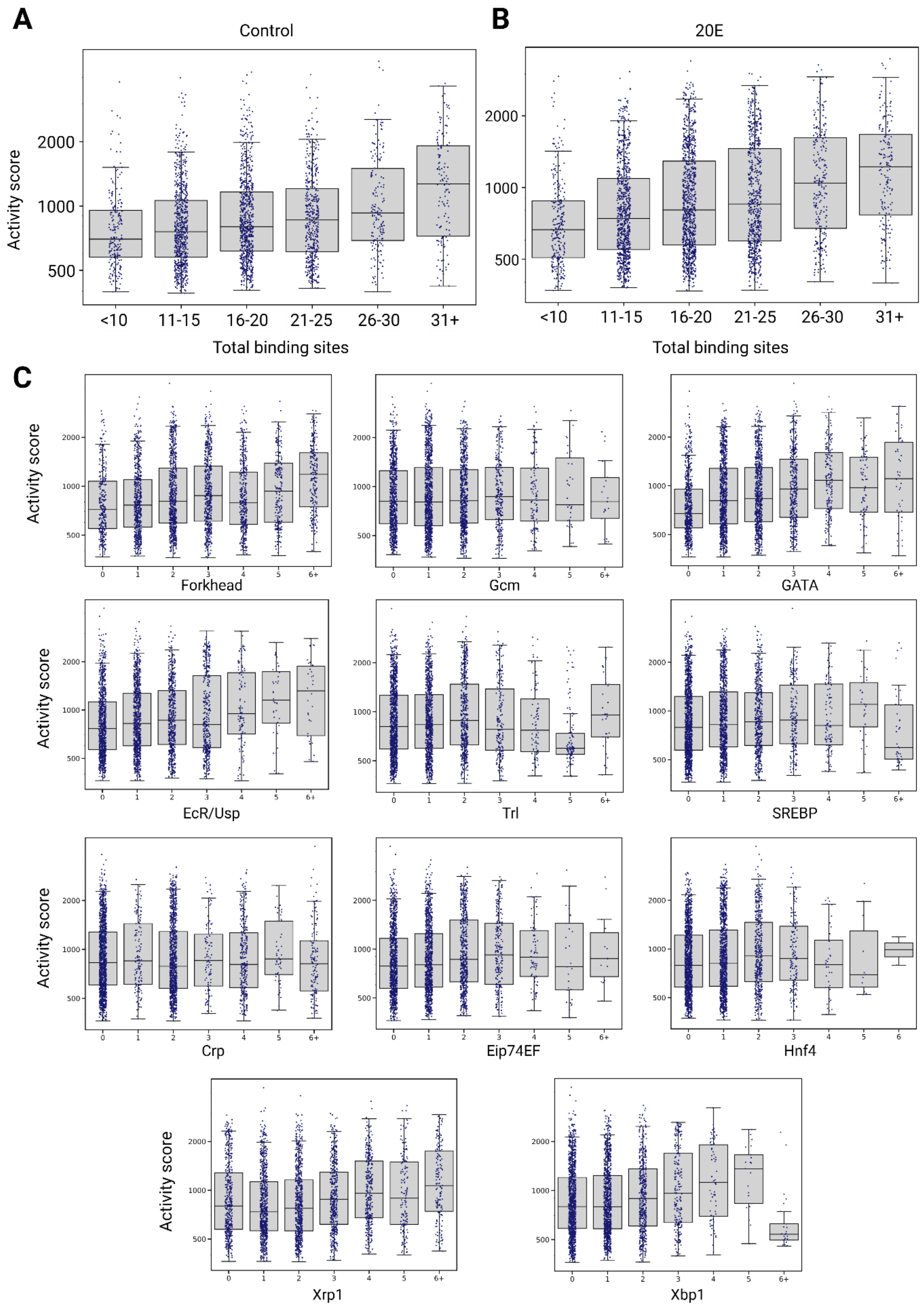


**Supplementary Figure S2: Activity score varies by number of TFBS.** A. Box and whiskers plot of STARR-seq activity score by total binding sites from 13 TFs in Control enhancers (A) and 20E (B) enhancers. (C) Activity scores for IMD enhancers by number of binding sites of individual TFs

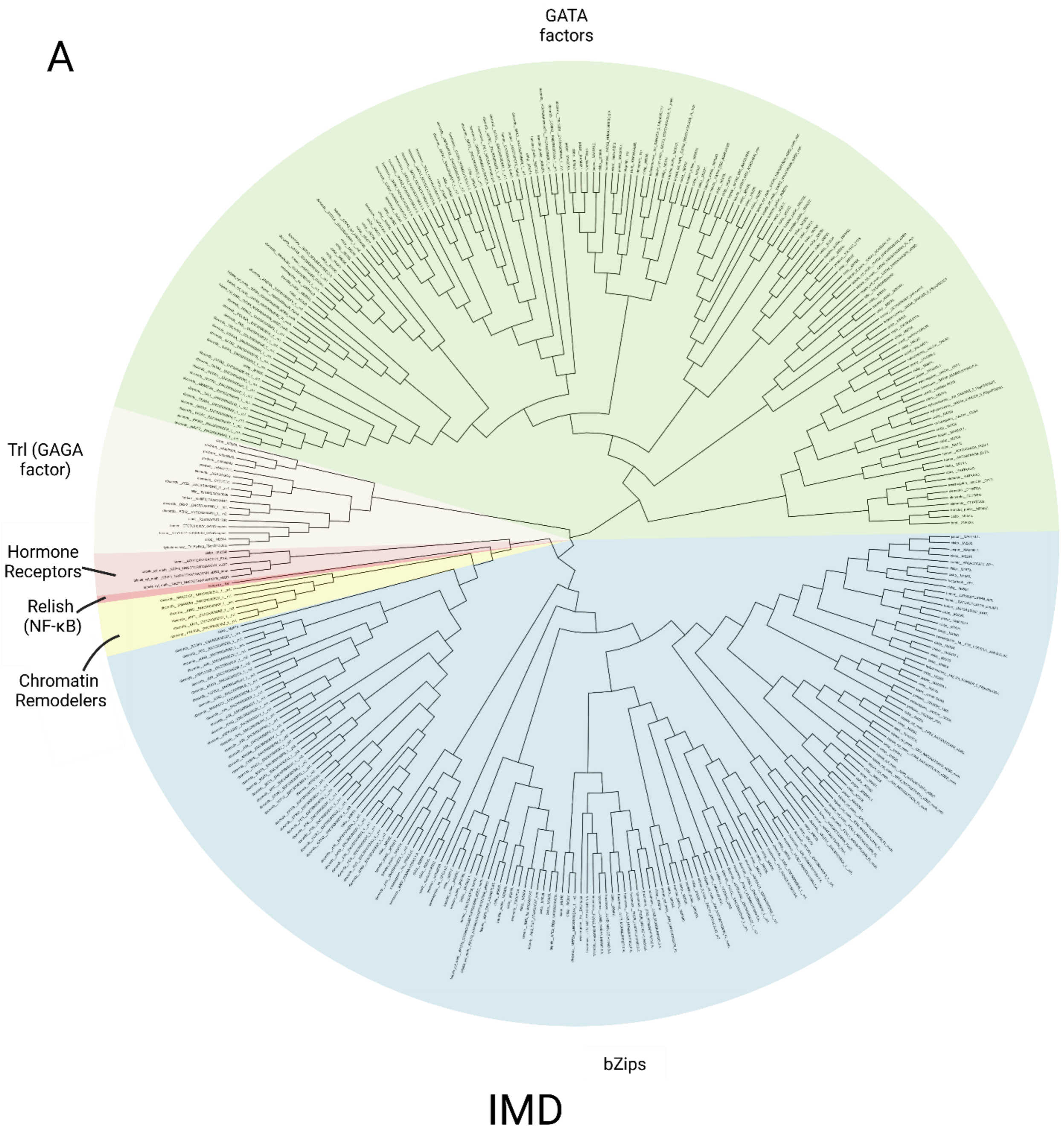
A
GATA factors
Trl (GAGA factor)
Hormone Receptors
Relish (NF-κB)
Chromatin Remodelers
bZips
IMD

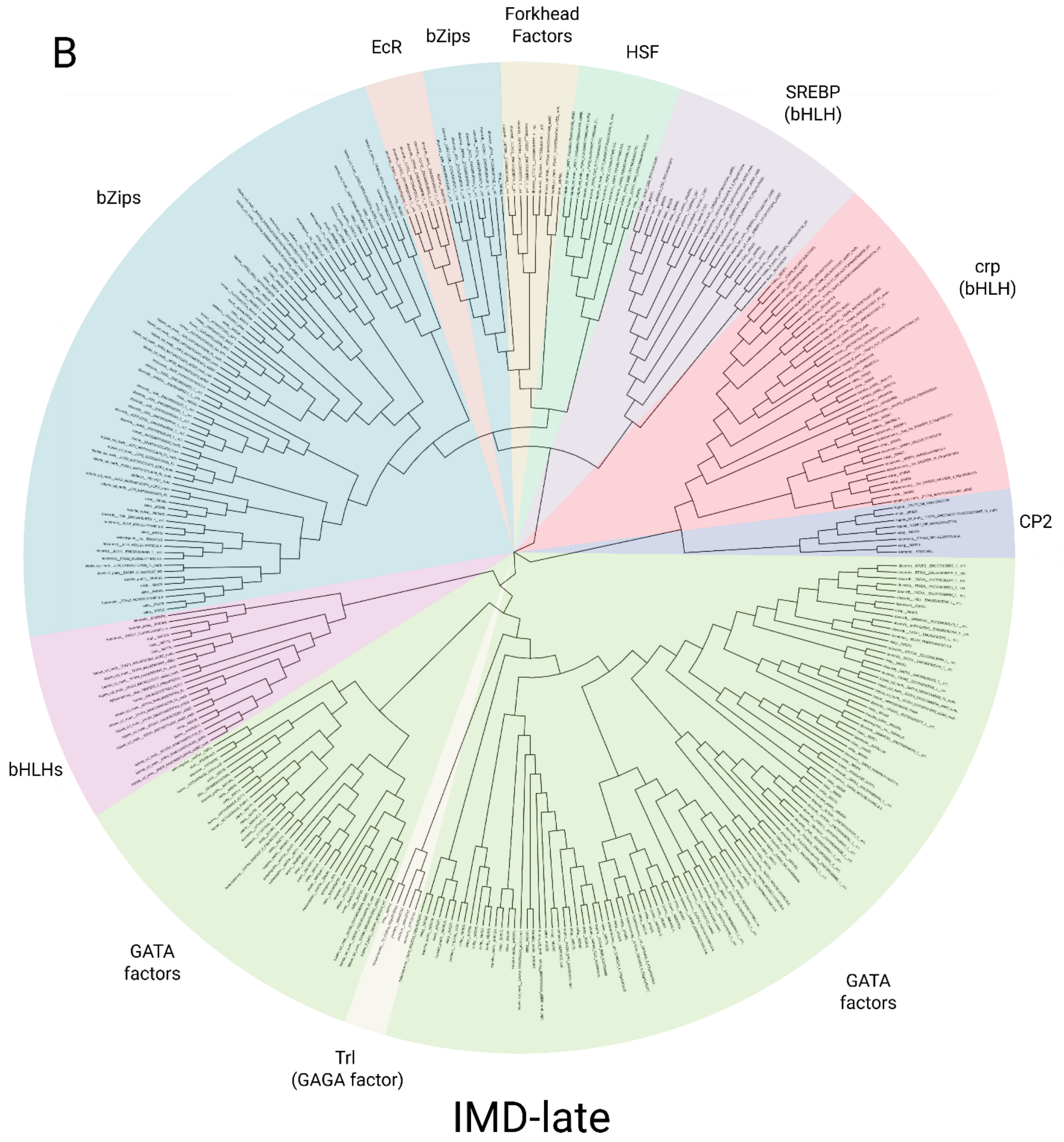


**Supplementary Figure S3: Several TF families are enriched in IMD enhancers.** (A) A motif tree of 360 motifs from icis-target analysis enriched three-fold in all IMD enhancers compared to the genome. TF families are indicated with colored sections. (B) A motif tree of 363 motifs from icis-target analysis enriched three-fold in IMD enhancers assigned to late acting genes.

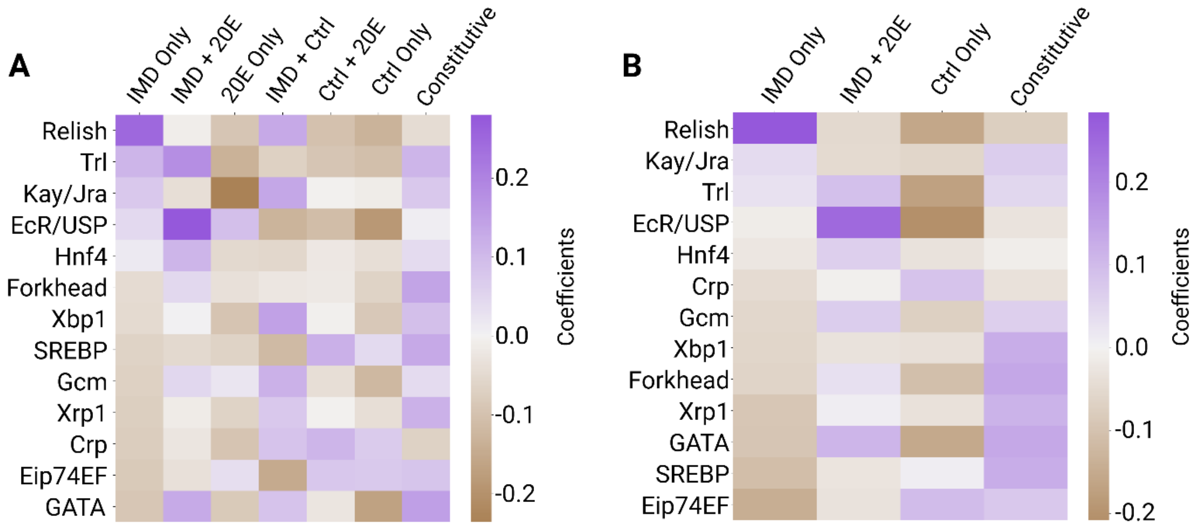


**C**

Predicted Classifications (columns); True Classifications (rows)

| | IMD Only | IMD + 20E | 20E Only | IMD + Ctrl | Ctrl + 20E | Ctrl Only | Constitutive |
|---|---|---|---|---|---|---|---|
| IMD Only | **27** | 7 | 15 | 15 | 3 | 10 | 3 |
| IMD + 20E | 21 | **57** | 34 | 28 | 17 | 20 | 28 |
| 20E Only | 14 | 8 | **27** | 7 | 4 | 15 | 9 |
| IMD + Ctrl | 3 | 0 | 1 | **3** | 3 | 5 | 3 |
| Ctrl + 20E | 6 | 5 | 6 | 6 | **2** | 8 | 6 |
| Ctrl Only | 17 | 10 | 29 | 23 | 10 | **39** | 11 |
| Constitutive | 24 | 50 | 42 | 46 | 18 | 31 | **78** |

**D**

Predicted Classifications (columns); True Classifications (rows)

| | IMD Only | IMD + 20E | Ctrl Only | Constitutive |
|---|---|---|---|---|
| IMD Only | **6** | 23 | 18 | 21 |
| IMD + 20E | 1 | **97** | 23 | 95 |
| Ctrl Only | 3 | 31 | **54** | 57 |
| Constitutive | 3 | 72 | 46 | **164** |

**Supplementary Figure S4: Logistic Regression analysis highlights the role of some TFs in predicting activity class.** (A) Coefficients of TF motifs from logistic regression classifying enhancers by activity class. (B) Coefficients from logistic regression classifying enhancers in four most biologically relevant activity classes. Positive coefficients are shown in purple, while negative coefficients are shown in brown. (C) Confusion matrix of logistic regression with a 20% test set size. Correctly called activity classifications are bolded and shaded in green. (D) Confusion matrix of four class logistic regression with a 20% test set size. Correctly called activity classifications are bolded and shaded in green.

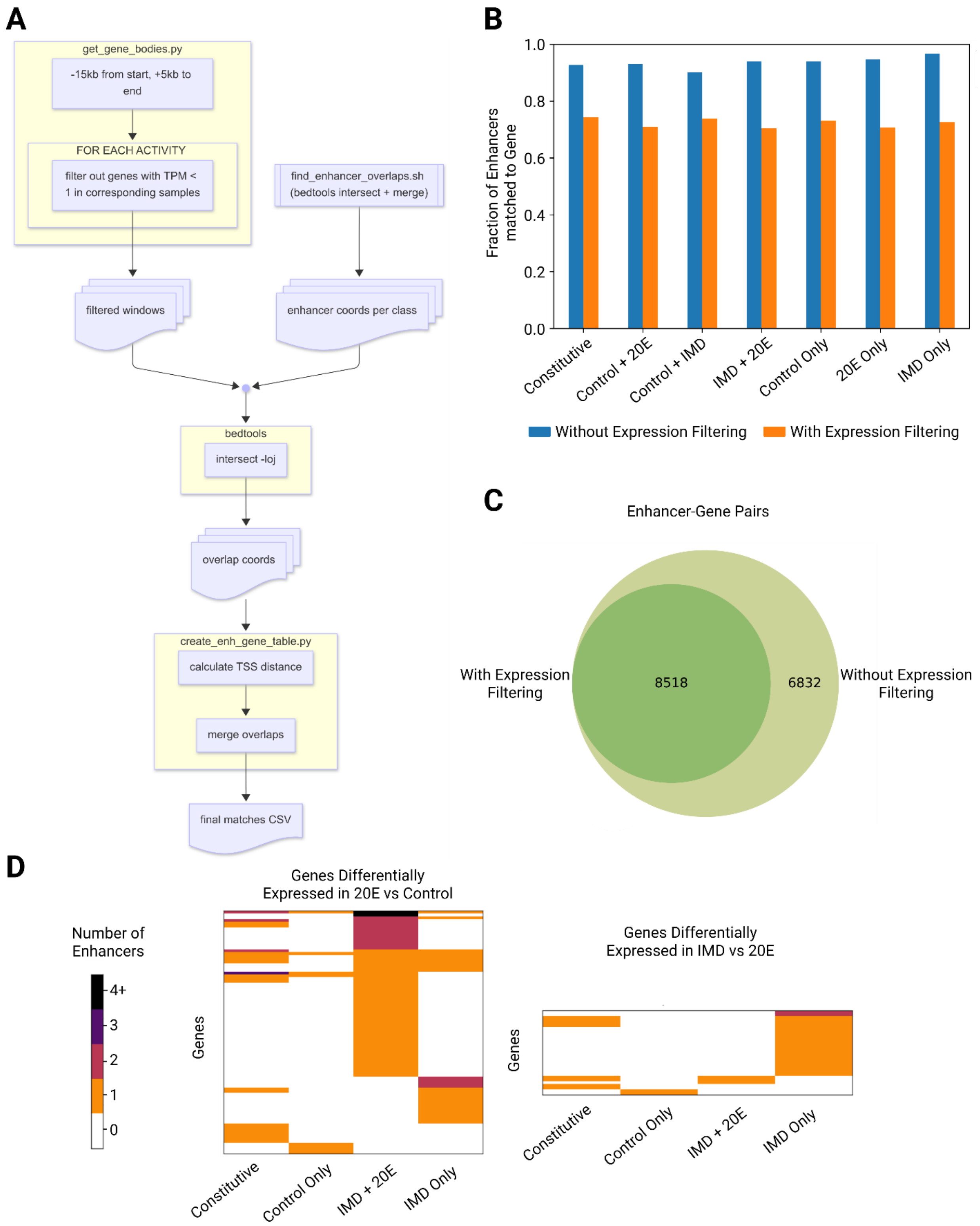


**Supplementary Figure S5: Expression filtering improves enhancer-gene assignments (A) Overview of gene-enhancer matching workflow.** (B) Fraction of enhancers matched to genes with and without expression filtering for all seven activity classes. (C) Venn Diagram of the total number of enhancer-gene pairs with and without expression filtering. Expression filtering removes a little less than half of the enhancer-gene pairs. (D) Enhancers assigned to genes upregulated (RNA-seq, log2 fold change > 2) in 20E vs Control (88 genes) and IMD vs 20E (31 genes) by activity class. Each row represents one gene and the number of enhancers in each activity class are indicated by the cell color.

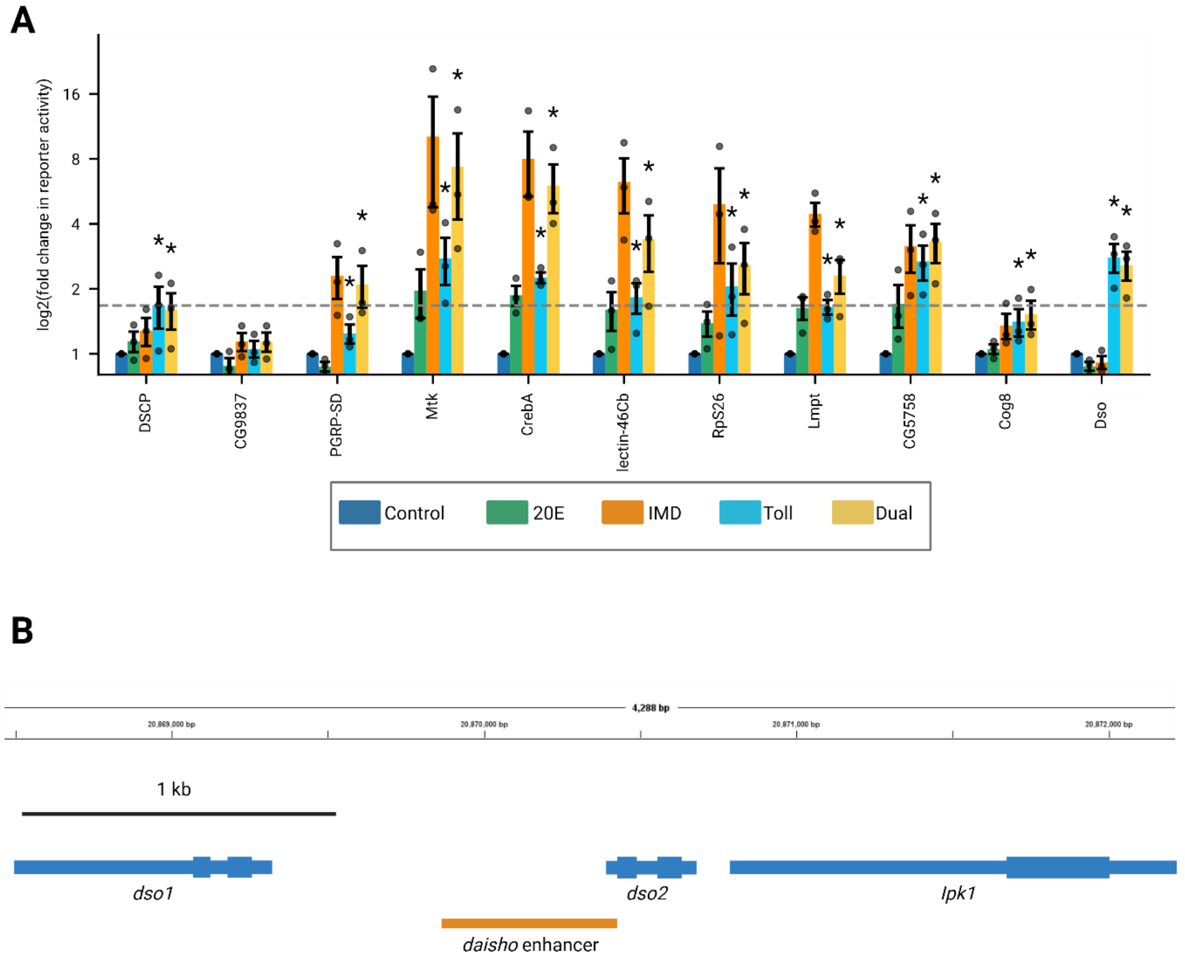


**Supplementary Figure S6: Enhancer reporters are differentially responsive to 20E, IMD, Toll and Dual Induction.** (A) The $\log_2$(fold change) of each reporter's activity, normalized to the control condition, was measured by flow cytometry. The bar shows the mean $\log_2$(fold change), the dots represent three biological replicates, and the error bars are the standard error of the mean. Each replicate measurement included at least 10,000 cells. Stars indicate a significant increase in activity between the control and starred conditions (one-sided Mann-Whitney U test; $p<0.05$). The dashed horizontal line is the mean fold change of the DSCP control construct in the Toll condition. The values for the 20E and IMD conditions for all reporters and the Toll and Dual conditions for the *daisho* reporter are also displayed in Figure 5, but are replicated here for ease of comparison to other conditions. (B) Genomic locus surrounding *daisho* enhancer. Genes are represented in blue, while the *daisho* STARR-seq enhancer is shown in orange.

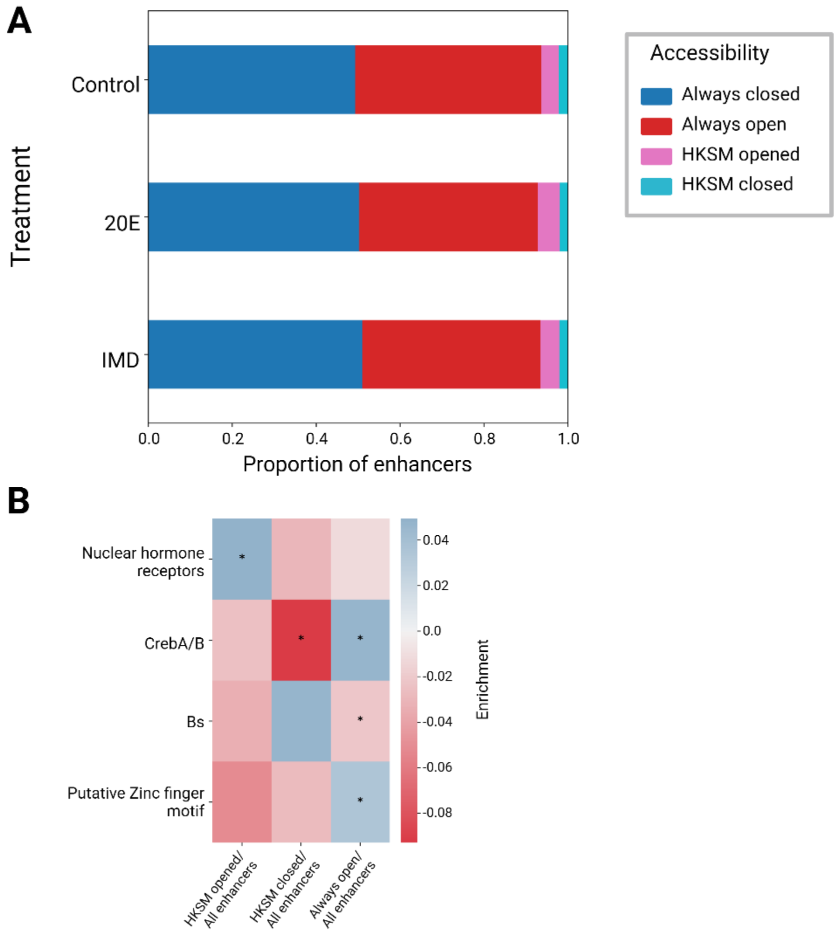


**Supplementary Figure S7: Most enhancers maintain chromatin structure upon immune stimulation.** (A) Fraction of enhancers of each accessibility group by treatment groups. (B) Log odds ratio of motifs enriched in accessibility grouped enhancers. We identified four distinct motifs: 1. A motif (GGTCA) assigned to several nuclear hormone receptors 2. A motif for either CrebA or CrebB. CrebA has been shown to act downstream of the Toll and IMD pathways to regulate tolerance to infection (Troha et al. 2018). 3. A motif for Blistered, known to play a role in wing development. 4.“GGTGTGTAT”, a highly enriched motif in HKSM opened enhancers compared to the genome. While no *Drosophila* TF was a clear match to this motif, we found the mouse C2H2 zinc finger Prdm14 had the closest known motif. We have called this sequence a putative zinc finger motif. Positive enrichment values are in blue and negative enrichment values are in red, and significance is indicated by * (p <0.05, Fisher’s test).

**Supplementary Table 1: Enhancer reporters**

| **Enhancer Name** | **Coordinates (dm6)** | **Mutated?** |
|---|---|---|
| *PGRP-SD* | 3L:7650744-7651565 | start codon mutated to ATT |
| *Cog8* | 2L:11445934-11446855 | no |
| *Mtk* | 2R:15408446-15409027 | start codon mutated to ATA |
| *CG5758* | 2L:18051506-18052253 | no |
| *Lmpt* | 3L:16878911-16879751 | no |
| *CrebA* | 3L:15539043-15539751 | no |
| *dso* | 2R:20869864-20870426 | no |
| *CG9837* | 3R:8813425-8814751 | no |
| *lectin-46Cb* | 2R:9808943-9809693 | no |
| *RpS26* | 2L:18335434-18336088 | no |
